# The nuclear PIWI-interacting protein Gtsf1 controls the selective degradation of small RNAs in *Paramecium*

**DOI:** 10.1101/2023.09.19.558372

**Authors:** Olivia Charmant, Julita Gruchota, Olivier Arnaiz, Coralie Zangarelli, Mireille Bétermier, Katarzyna Nowak, Véronique Legros, Guillaume Chevreux, Jacek Nowak, Sandra Duharcourt

## Abstract

Ciliates undergo developmentally programmed genome elimination, in which small RNAs direct the removal of target DNA segments, including transposable elements. At each sexual generation, the development of the macronucleus (MAC) requires massive and reproducible elimination of a large proportion of the germline micronuclear (MIC) genome, leading to a highly streamlined somatic MAC genome. 25-nt long scnRNAs are produced from the entire germline MIC genome during meiosis, and this initial complex small RNA population is then transported to the maternal MAC, where selection of scnRNAs corresponding to germline (MIC)-specific sequences is thought to take place. Selected scnRNAs, loaded onto the PIWI protein Ptiwi09, guide the deposition of histone H3 post-translational modifications (H3K9me3 and H3K27me3) onto transposable elements in the developing macronucleus, ultimately triggering their specific elimination. How germline-specific MIC scnRNAs are selected remains to be determined. Here, we provide important mechanistic insights into the scnRNA selection pathway by identifying a *Paramecium* homolog of Gametocyte specific factor 1 (Gtsf1) as essential for the selective degradation of scnRNAs corresponding to retained somatic MAC sequences. Consistently, we also show that Gtsf1 is exclusively localized in the maternal macronucleus and associates with the scnRNA-binding protein Ptiwi09. Furthermore, Gtsf1 is necessary for DNA elimination and correct H3K9me3 and H3K27me3 localization in the new developing macronucleus, demonstrating that the scnRNA selection process is important for genome elimination. We propose that Gtsf1 is required for the coordinated degradation of Ptiwi09-scnRNA complexes that pair with nascent RNA transcribed from the maternal MAC genome, similarly to the mechanism suggested for microRNA target-directed degradation in metazoans.

## Introduction

One major host defense mechanism to silence transposable elements (TE) is the small RNA silencing pathway. Small RNAs of 20-30 nt in length are loaded onto PIWI proteins to form small RNA-PIWI silencing complexes, which pair with nascent transcript by sequence complementarity, recruit histone methyltransferases to chromatin and repress the transcriptional activity of TEs and other repeats (Haase, 2022). Small RNA sequences are highly diverse and yet highly specific to TEs. The mechanisms underlying the distinction between TEs and the rest of the genome during the establishment of small RNA populations remain to be fully understood.

In animal gonadal cells, such as the *Drosophila* ovaries, Piwi-interacting RNAs (piRNAs) are mostly derived from heterochromatic TE-rich loci known as the piRNA clusters, and cluster transcripts must be specifically selected for piRNA biogenesis (Czech et al., 2018). Thus, the vast majority of these small RNAs originate from distinct genomic hotspots. The logic is radically different in the ciliate *Paramecium*, in which the entire germline genome initially produces 25-nt scnRNAs from both TE and non-TE sequences (Lepere et al., 2009; Miró-Pina et al., 2023; Sandoval et al., 2014; Singh et al., 2014). In subsequent steps, the subpopulation of scnRNAs that correspond to TEs is selected to trigger the physical removal of TEs from the genome, a definitive form of TE silencing (Bouhouche et al., 2011; Furrer et al., 2017; Lepere et al., 2009, 2008). The mechanism that enables the specific selection of scnRNAs corresponding to TEs is currently unknown.

In *Paramecium*, the germline and somatic functions are supported by two types of nuclei that coexist in the same cytoplasm. The diploid germline micronucleus (MIC) transmits the genetic information from one generation to the next, while the polyploid somatic macronucleus (MAC) ensures gene expression, but is destroyed at each sexual cycle. During the self-fertilization process of autogamy, the MIC undergoes meiosis and karyogamy to produce the zygotic nucleus. New MICs and new MACs develop from mitotic products of the zygotic nucleus. During development of the new MAC, massive and reproducible elimination of nearly 30% of the genome (∼30 Mb out of 108 Mb) of specific germline sequences occurs (Betermier and Duharcourt, 2014). Eliminated germline sequences include 45,000 Internal Eliminated Sequences (IESs) that are remnants of TEs scattered throughout the genome (Arnaiz et al., 2012; Sellis et al., 2021). Other Eliminated Sequences (OES) (Swart et al., 2017) correspond to large regions comprising repeats such as TEs and satellites (Guérin et al., 2017). Understanding how such diverse sequences are defined and eliminated despite the lack of conserved sequence motifs remains challenging.

Previous studies have shown that the specific recognition of TEs destined for elimination in fact involves scnRNAs which direct histone mark deposition and subsequent DNA excision in the developing MAC genome (Bischerour et al., 2018; Miró-Pina et al., 2022). In a mechanism very similar to transcriptional TE silencing described in fungi and metazoans (Martienssen and Moazed, 2015), scnRNAs target the Piwi proteins to complementary nascent transcripts (Maliszewska-Olejniczak et al., 2015) to guide histone modifications on TE loci in the new MAC (Frapporti et al., 2019), thereby providing the required sequence specificity. A physical interaction between the scnRNA-binding protein Ptiwi09 and the Polycomb Repressive Complex 2 (PRC2-Ezl1) was recently shown to mediate the establishment of histone H3K9me3 and H3K27me3 at scnRNA-targeted regions (Miró-Pina et al., 2022; Wang et al., 2022). Deposition of these repressive chromatin marks in the new MAC is essential for the elimination of TEs and of 70% of IESs (Frapporti et al., 2019; Lhuillier-Akakpo et al., 2014; Miró-Pina et al., 2022).

25-nt long scnRNAs are produced during meiosis from the whole germline MIC genome, which is entirely transcribed by RNA polymerase II with the participation of a specialized Spt5/Spt4 elongation complex (Gruchota et al., 2017; Khurana et al., 2014; Mochizuki and Gorovsky, 2004; Owsian et al., 2022). scnRNA biogenesis relies on a developmental-specific RNA interference (RNAi) pathway that involves the Dicer-like proteins Dcl2 and Dcl3 (Lepere et al., 2009; Miró-Pina et al., 2023; Owsian et al., 2022; Sandoval et al., 2014; Singh et al., 2014). These scnRNAs then bind to Ptiwi01/09 proteins and are transported to the maternal MAC (Furrer et al., 2017), where selection of scnRNAs is believed to occur. Working models posit that scnRNAs are sorted out by pairing interactions with nascent non-coding RNAs produced by the MAC genome (Aronica et al., 2008; Lepere et al., 2008; Mochizuki et al., 2002). Non-coding RNAs are thought to trigger the degradation of their cognate small RNA (MAC-scnRNAs), while allowing scnRNAs corresponding to MIC-specific sequences (thereafter called MIC-scnRNAs), which by definition cannot pair with MAC transcripts, to be retained. This selective degradation of MAC-scnRNAs would thus result in the specific selection of the subpopulation corresponding to MIC-scnRNAs. The requirement of complementary maternal MAC transcripts for scnRNA selection has been directly demonstrated (Lepere et al., 2008). On the other hand, how MAC-scnRNAs are degraded and how scnRNAs corresponding to TEs are selected is currently unknown.

Approximately 60% of the scnRNA population (representing the 72 Mb corresponding to the MAC genome) is degraded, making *Paramecium* an exquisite model to decipher the underlying molecular mechanisms of this selective degradation. To uncover candidate proteins involved in the process, we sought to identify the protein partners of the scnRNA-binding protein Ptiwi09 during the developmental stage at which scnRNA selection is thought to occur. We discover that *Paramecium* Gtsf1 is a nuclear Ptiwi09-interacting protein in the maternal MAC. The conserved Asterix/Gametocyte-specific factor 1 (GTSF1) proteins are essential piRNA factors that contribute to the repression of transposons in mice, *Drosophila* and zebrafish (Ipsaro and Joshua-Tor, 2022). We show that *Paramecium* Gtsf1 is involved in scnRNA-guided TE repression by controlling the degradation of MAC-scnRNAs and of the Ptiwi09 protein. We further demonstrate that defective scnRNA degradation leads to histone modification and scnRNA-guided DNA elimination defects. We propose that Gtsf1 is required for the coordinated degradation of the Ptiwi09 proteins and the bound scnRNAs when engaged in paring interactions with complementary nascent maternal MAC RNA, similarly to the proposed mechanism for microRNA target-directed degradation.

## Results

### Identification of Gtsf1, a Ptiwi09-interacting protein in the maternal MAC

To uncover candidate proteins involved in scnRNA selection, we identified the protein partners of Ptiwi09 during the developmental stage at which scnRNA selection occurs. Whole cell extracts were prepared from *Paramecium* control cells and cells expressing a 3xFLAG-tagged Ptiwi09, at an early stage of the sexual process of autogamy, when the protein is present in the maternal MAC (Figure 1A, Figure S1). Immunoprecipitation (IP) of the FLAG tag, in 2 and 4 replicates for the control and Ptiwi09, respectively (Methods), was followed by mass spectrometry (MS) (Figure 1B and 1C). Statistical analyses revealed 283 differential proteins (out of 633 identified proteins) in the Ptiwi09 IP compared with control (fold change > 2; p value < 0.05; unique peptide > 2) (Figure 1C, Table S1). We recovered Ptiwi09 as expected, and its paralog Ptiwi03, a previously reported partner of Ptiwi09 in the new developing MAC (Figure 1C; (Miró-Pina et al., 2022)). We also identified PRC2 core components (Ezl1, Caf1, Eed) and PRC2 cofactors (Rf2 and Rf4) (Figure 1C, Table S1). These proteins were recently shown to be involved in the scnRNA selection process (Miró-Pina et al., 2023; Wang et al., 2022), and the Rf4 cofactor was further shown to physically interact with Ptiwi09 in the new developing MAC (Miró-Pina et al., 2022). Another Ptiwi09 interactor we identified was an uncharacterized, small protein (16 kDa) referred to as Gametocyte specific factor 1 (Gtsf1) (Figure 1C, Figure 2A, Table S1). Interestingly, it had not been identified among the protein partners of Ptiwi09 when purified from the new developing MAC at a later stage of autogamy (Miró-Pina et al., 2022). Like its counterparts in other organisms *(D. melanogaster* (Dönertas et al., 2013)*, M. musculus* (Yoshimura et al., 2009), and *C. elegans* (Almeida et al., 2018)), *Paramecium* Gtsf1 contains a double CHHC zinc finger domain at the N-terminus and an unstructured C-terminus (Figure 2A). RNA-seq expression data (Arnaiz et al., 2017) shows that the *GTSF1* gene displays a developmental expression profile similar to that of *PTIWI09* and *EZL1* (Figure 2B). To investigate Gtsf1 function, we performed KD experiments using RNAi during the sexual cycle of autogamy. KD of *GTSF1* leads to lethality of the sexual progeny, whereas KD of a control non-essential gene leads to survival of the post-autogamous progeny (Figure 2C, Table S2). Thus, *GTSF1* is an essential gene during the sexual cycle.

**Figure 1.**
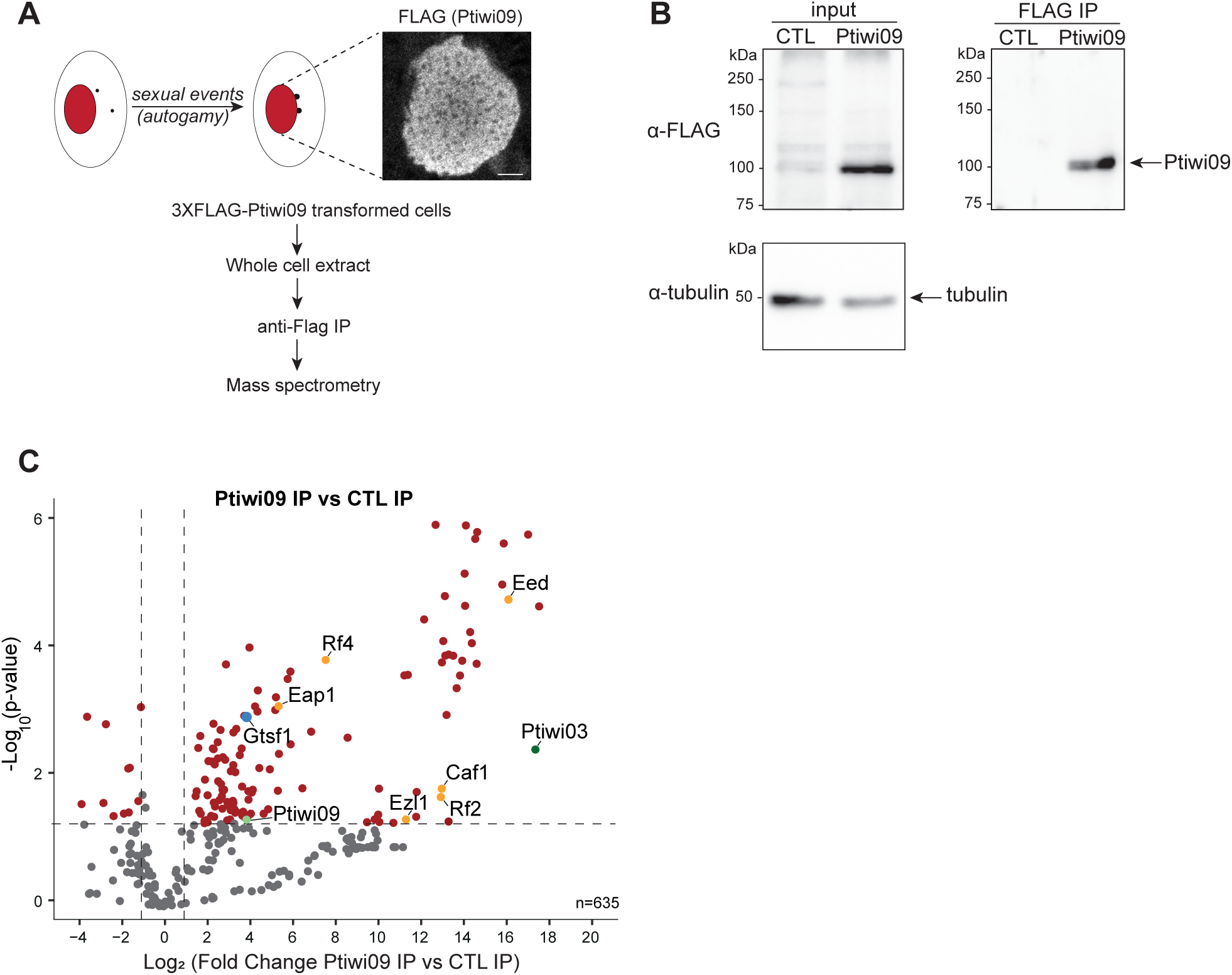
Identification of Ptiwi09-interacting partners in the maternal macronucleus. A. Schematic representation of Ptiwi09 immunoprecipitation experiments. Anti-FLAG immunostaining of cells transformed with a *3xFLAG-PTIWI09* transgene when whole cell extracts for immunoprecipitation were performed (T=0 hours, at the onset of autogamy). Scale bar, 5 μm. B. Western blot analysis of whole cell extracts from cells expressing *3XFLAG-PTIWI09* (Ptiwi09) and non-injected cells (CTL) before (input) or after affinity purification (IP). FLAG antibodies were used for 3XFLAG-Ptiwi09 detection and tubulin antibodies for normalization. C. Volcano plot of the quantitative label-free mass spectrometry analysis of 3XFLAG-Ptiwi09 affinity purification. Significantly enriched proteins in Ptiwi09 IP (4 replicates) over control IP (2 replicates) (the fold change is greater than 2, the total number of peptides is greater than 2 and the p-value is less than 0.05) are in the upper right corner. The bait Ptiwi09 (light green), Gtsf1 (blue), PRC2-Ezl1 complex (orange) and Ptiwi09-inte-racting proteins (dark green) are highlighted.

**Figure 2.**
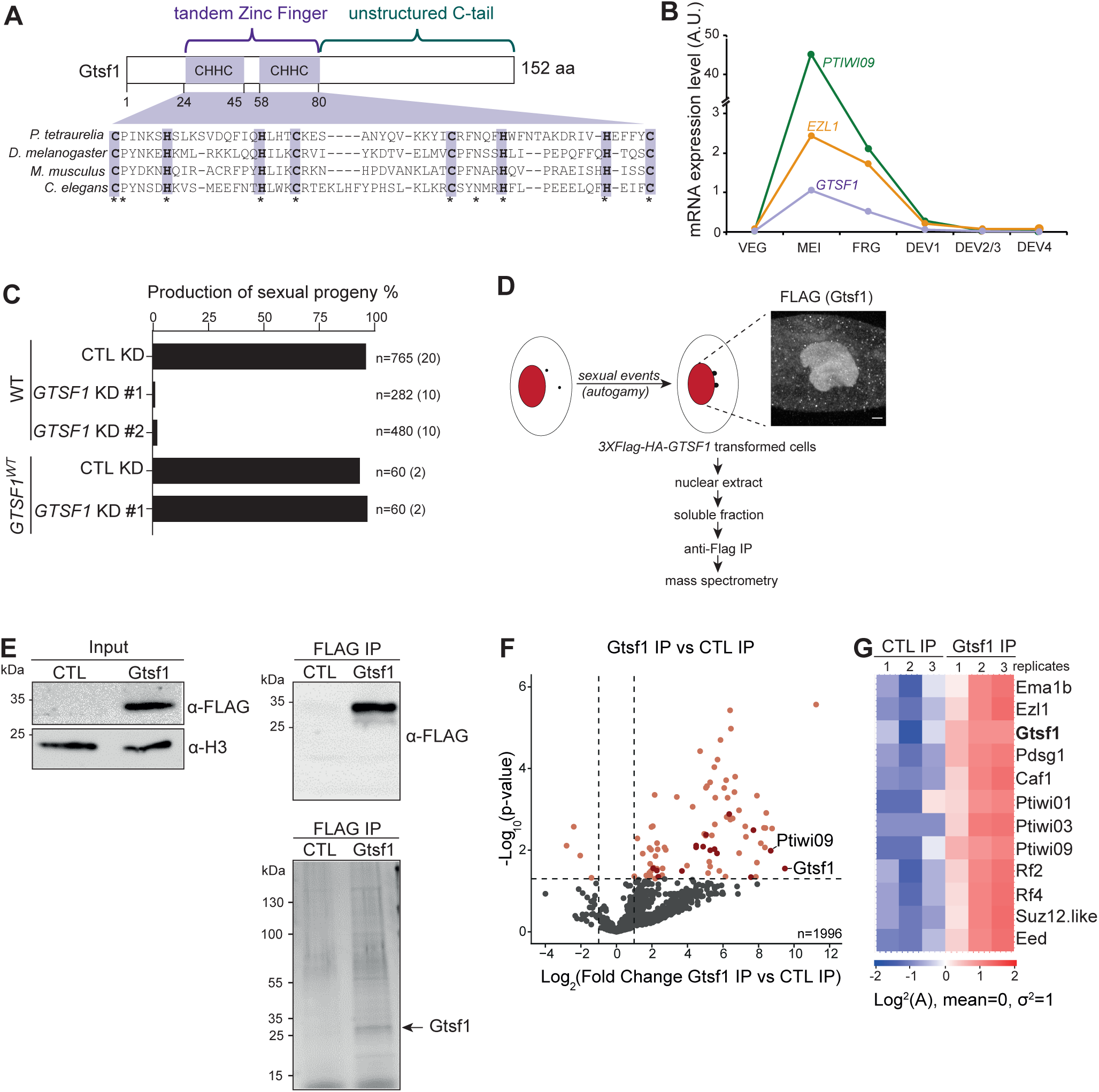
Gtsf1 is an essential nuclear protein that interacts with Ptiwi09. A. Schematic representation of Gtsf1-predicted protein domains and alignment of the N-terminal portion of Gtsf1 from *P. tetraurelia* (PTET.51.1.G0490019), *D. melanogaster* (CG3893), *M. musculus* (MGI:1921424) and *C. elegans* (CELE_T06A10.3) (from top to bottom). The conserved tandem CHHC zinc finger domain is highlighted in purple. B. mRNA expression levels of the genes encoding *GTSF1* (purple), *EZL1* (yellow) and *PTIWI09* (green) at different developmental stages during autogamy in arbitrary units (Arnaiz et al., 2017). C. Production of sexual progeny of wild-type cells (WT) and cells expressing a 3XFLAG-HA-Gtsf1 (*GTSF1*^WT^) fusion protein following *GTSF1* (with 2 RNAi inserts #1 and #2) or *ICL7* (CTL) RNAi-mediated silencing (KD). The total number of cells analyzed for each RNAi and the number of independent experiments (in parentheses) are indicated. D. Schematic representation of Gtsf1 immunoprecipitation experiments. Anti-FLAG immunostaining of cells transformed with a *3XFLAG-HA-GTSF1* transgene, at the same time point as that used for preparation of nuclear extracts for immunoprecipitation (T=0 hours). Scale bar, 10 μm. E. Top: Western blot analysis of nuclear extracts of *Paramecium* expressing a 3XFLAG-HA-Gtsf1 functional protein (Gtsf1) or 3XFLAG-HA (CTL) before (input) or after affinity purification (FLAG IP). Bottom: silver-stained gel of pulled-down proteins. Predicted 3XFLAG-HA-Gtsf1 MW: 23.6kDa. F. Volcano plot of the quantitative label-free mass-spectrometry analysis of 3xFLAG-HA-Gtsf1 affinity purification. 3 replicates were analyzed for each condition. Significantly enriched proteins in Gtsf1 IP over control IP (the fold change is greater than 2, the total number of peptides is greater than 2 and the p-value is less than 0.05) that are encoded by developmental genes with an early expression peak are highlighted in dark red, and include Gtsf1 and Ptiwi09. G. Heatmap of abundance of the 12 top protein hits in Gtsf1 IP over control IP. Log2 of abundance is transformed and standardized with a mean of 0 and variance of 1. Only the proteins encoded by developmental genes with an early expression peak were considered.

To confirm the interaction between Gtsf1 and Ptiwi09, we performed reciprocal IPs using a functional tagged 3XFLAG-HA-Gtsf1 protein capable of rescuing the lethality caused by *GTSF1* KD (Figure 2C). To immunoprecipitate Gtsf1, we prepared nuclear extracts from control *Paramecium* cells and cells expressing the functional fusion protein. As expected, immunofluorescence experiments confirmed that Gtsf1, like Ptiwi09 (Bouhouche et al., 2011; Miró-Pina et al., 2023), localizes in the maternal macronucleus during meiosis (Figure 2D). Immunoprecipitation (IP) of the FLAG tag in triplicates for control and Gtsf1 (Figure 2E, Figure S1) was followed by MS (Figure 2F and 2G). MS analyses revealed that Gtsf1 (bait) and the scnRNA-binding protein Ptiwi09 were significantly enriched in Gtsf1 IPs compared with controls, confirming the interaction between Gtsf1 and Ptiwi09 (Figure 2F). Statistical analyses revealed 95 differential proteins (95 of 1996 identified proteins) in the Gtsf1 IP compared with control (fold change > 2; p value < 0.05; unique peptides > 2) (Figure 2F, Table S1).The top differential proteins encoded by developmental genes with an early expression peak, similar to Gtsf1 and Ptiwi09, included the Ptiwi09 paralogs Ptiwi01 and Ptiwi03 (Bouhouche et al., 2011) and the Ptiwi09-associated protein Ema1b (Miró-Pina et al., 2022), a putative RNA helicase involved in DNA elimination (Nowak et al., 2011) (Figure 2F and 2G, Table S1). Its *Tetrahymena* counterpart (Ema1p) was reported to be required for scnRNA selection (Aronica et al., 2008). We also identified the PRC2 core components Ezl1, Caf1, Suz12-like and Eed, and PRC2 cofactors Rf2 and Rf4, confirming the interaction between Gtsf1, Ptiwi09 and PRC2. In addition, we identified Pdsg1, a protein previously reported to be involved in IES elimination and scnRNA selection (Arambasic et al., 2014).

To gain further insight into the role of Gtsf1, we examined its subcellular localization. Co-immunofluorescence experiments were performed with *Paramecium* cells expressing the N-terminally FLAG-tagged Gtsf1 under its endogenous regulatory regions at different stages of sexual events using FLAG antibodies and specific H3K27me3 antibodies (Frapporti et al., 2019) (Figure 3). The FLAG signal is detected in the MAC at the onset of meiosis I, as seen with H3K27me3 labeling of the MICs (Lhuillier-Akakpo et al., 2014). Gtsf1 is detected in the maternal MAC before this nucleus is fragmented, and persists until the formation of the new developing MAC, when the signal vanishes from the fragmented MAC. Thus, Gtsf1 is the first protein identified so far that exclusively localizes in the maternal MAC during MIC meiosis, and is absent from the new MAC.

**Figure 3.**
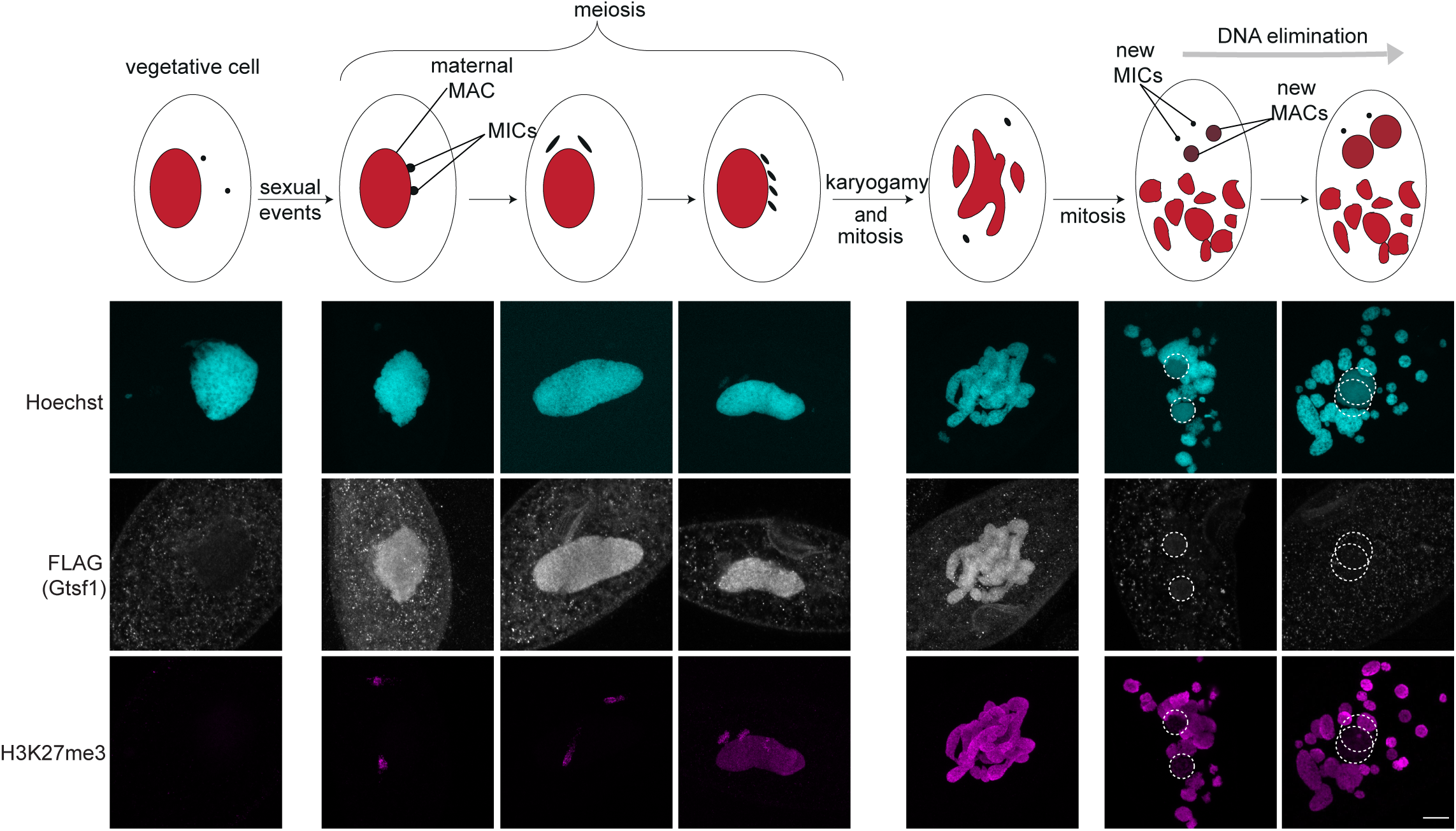
Gtsf1 is exclusively localized in the maternal MAC. Co-immunostaining with FLAG and H3K27me3 antibodies on cells expressing 3XFLAG-HA-Gtsf1 during vegetative life and at different stages of the *Paramecium* sexual cycle (autogamy). After MIC meiosis and karyogamy, the zygotic nucleus undergoes two successive mitoses producing four nuclei that differenciate into new MICs and new MACs. Representative images are displayed. Overlay of Z-projection of magnified views of Hoechst staining (top), FLAG- (middle) and H3K27me3-(bottom) specific antibodies are presented. Dashed white circles indicate the new developing MACs. The other Hoechst-stained nuclei are the fragments of the maternal MAC. Note that meiotic MICs are labelled with H3K27me3. Scale bar, 10 μm.

### Gtsf1 is required for efficient DNA elimination and TE silencing

Given that Gtsf1 is a Ptiwi09 partner and that depletion of the Ptiwi01/09 proteins impairs DNA elimination (Bouhouche et al., 2011; Furrer et al., 2017; Miró-Pina et al., 2022), we examined the impact of *GTSF1* KD on DNA elimination. Sequencing of genomic DNA extracted from two preparations enriched for new MACs upon depletion of Gtsf1 (*GTSF1^enriched^*) (Figure 4A; Figure S2) indicated that 3.3% to 6.6% of IESs are significantly retained (1499 and 3001 respectively, p-value < 0.05) (Figure 4B; Figure S2). The smaller subset was almost totally included in the larger one, with 3,285 affected IESs in both samples (Figure 4C). This number is likely underestimated because of contamination of the sample preparation by fragments of the maternal MAC. For this reason, we performed sequencing of genomic DNA extracted from new MACs sorted by flow cytometry from cells depleted for *GTSF1* (*GTSF1^sorted^*) (Figure 4A, Figure S3). We found that 35.7% of IESs, a larger subset indeed, are significantly retained (16098, p-value < 0.05) (Figure 4B, Figure S2). Previous studies showed that a small subset of IESs depend on Ptiwi01/09 (6.1%) (Furrer et al., 2017; Miró-Pina et al., 2022), all of which require Ezl1 for their excision (Lhuillier-Akakpo et al., 2014). We found that IESs retained in *GTSF1* KD are essentially all (97.3 %) included in the *EZL1*-dependent subset and that they are enriched in scnRNA-dependent IESs (Figure 4C-D; Figure S2). We also noted that Gtsf1-dependent IESs are enriched for longer IESs (Figure S2).

**Figure 4.**
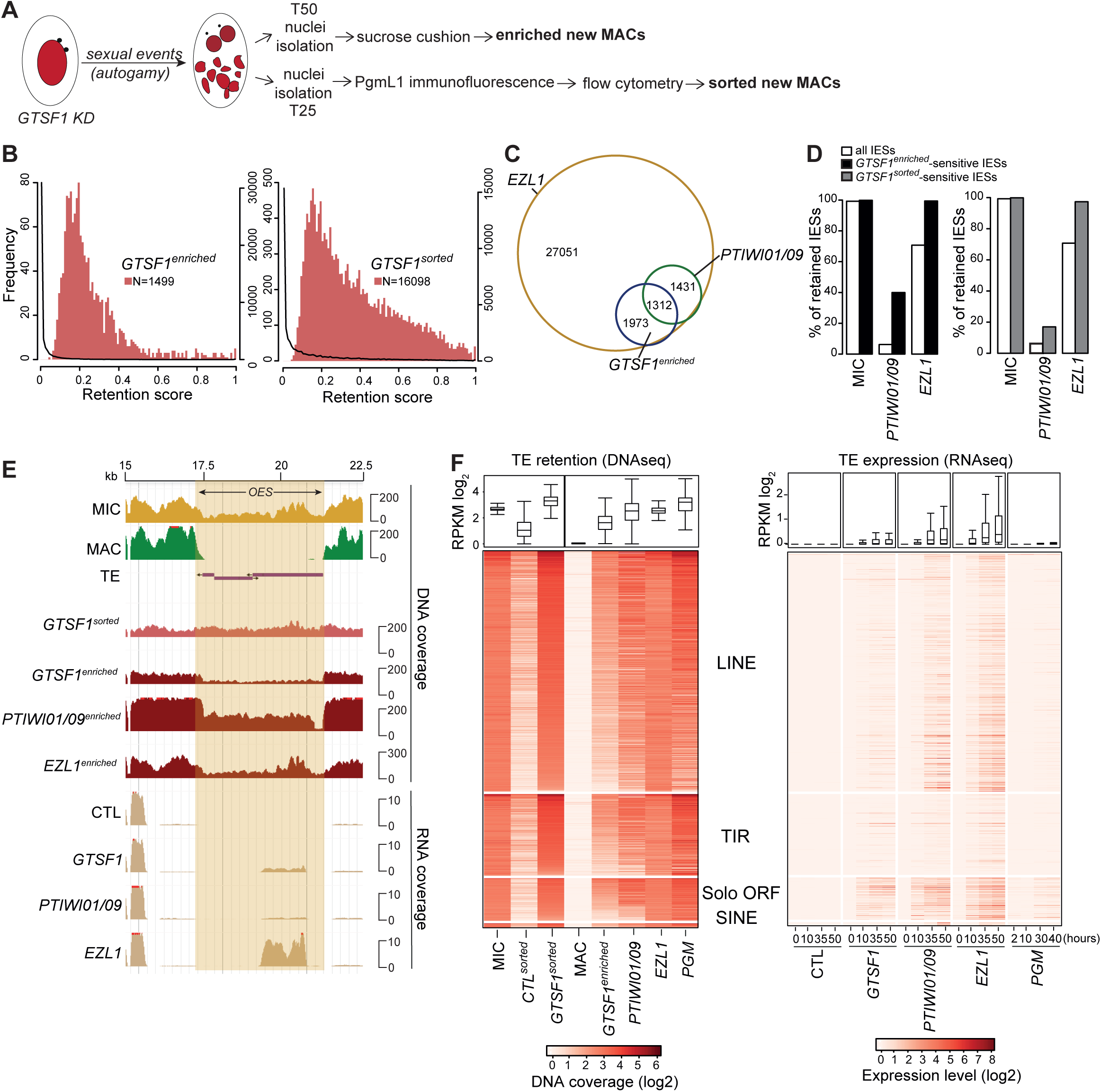
Gtsf1 is required for efficient DNA elimination and TE silencing. A. Schematic representation of the experimental design to extract genomic DNA from *GTSF1*-silenced cells. B. Histograms of IES retention scores upon *GTSF1* KD. The significantly retained IESs in *GTSF1* KD are represented by the red histograms (scale on the left), while the global distribution for all IESs retained in *GTSF1* KD is represented by the black curve (scale on the right). C. Venn diagram of significantly retained IESs upon different KDs (enriched new MACs in all conditions, 2 *GTSF1*-enriched replicates (R1 and R2) are combined, Figure S2). D. Histogram of the percentage of retained IESs (all (white), *GTSF1*-enriched (black), *GTSF1-*sorted (grey)) in MIC, *EZL1* or *PTIWI01/09* KDs. E. Representative genomic region depicting DNA coverage and RNA coverage (T= 50 hours) in *ND7* (CTL), *GTSF1*, *EZL1* and *PTIWI01/09* KD (NODE_36852_length_47919_cov_44.754879 between 15 and 22.5 kb). F. Left panel: Heatmaps of TE normalized DNA coverage in each KD. Right panel: Heatmap of RNA expression levels at different time points during development in each KD. TE copies are ordered by the mean DNA coverage of *GTSF1*-enriched and *GTSF1*-sorted in each family. The boxplots show the coverage (RPKM log2) for all TE copies.

To analyze the effects on the elimination of MIC-limited sequences other than IESs, we examined TEs and found that all four of the major distinct TE families (Guérin et al., 2017) are retained upon *GTSF1* KD, similarly to *EZL1* and *PTIWI01/09* KDs (Figure 4E-F; Figure S2). Given that TE transcript levels are increased upon depletion of Ptiwi01/09 and of the PRC2 components (Miró-Pina et al., 2022), and to determine whether this was also the case upon depletion of Gtsf1, we performed RNA-seq at the same developmental stages upon *GTSF1* KD (T=0, T= 10, T=35 and T=50 hours after the onset of autogamy) (Figure 4E-F; Figure S1). 5% of all annotated TE copies become expressed (>1 RPKM) during MAC development (T = 50 h) upon *GTSF1* KD (Figure 4E-F). TE de-silencing in Gtsf1-depleted cells is weaker than in Ptiwi01/09- (14%) or Ezl1-depleted cells (25%) but is specific, as no TE expression is detected in cells depleted for the elimination machinery (Pgm), when all DNA elimination events are blocked (Figure 4F).

In addition to TEs, the expression of a few thousand developmental genes is deregulated upon *EZL1* KD (Frapporti et al., 2019), raising the question of whether Gtsf1 might contribute to their regulation as well. We therefore used our RNAseq dataset to evaluate the impact of *GTSF1* KD on the expression of genes whose expression is either up-(N=1505) or down-(N=870) regulated upon *EZL1* KD. We found that these genes are indeed deregulated upon *GTSF1* KD (Figure S2). Focusing on the 628 developmental genes that are up-regulated upon KD of the elimination machinery (Pgm, Ku80 and Xrcc4) (Bazin-Gélis et al., 2023), we found that depletion of either Gtsf1 or Ezl1 also affects the expression of these genes (Figure S2). Thus, Gtsf1 depletion, as that of other proteins essential for DNA elimination, results in the deregulation of a subset of developmental genes, and this is likely a response to defective IES excision in the new MAC.

### Gtsf1 depletion affects H3K9me3 and H3K27me3 levels and localization

We examined whether depletion of Gtsf1 had an effect on accumulation of H3K9 and H3K27 trimethylation during autogamy using immunofluorescence. H3K9me3 and H3K27me3 accumulate in the new MAC in control conditions (Figure 5A), as previously described (Frapporti et al., 2019; Lhuillier-Akakpo et al., 2014). In contrast, H3K9me3 and H3K27me3 no longer accumulate in the new MAC if any of the core components of PRC2 is absent, and their levels are diminished in the absence of Ptiwi01/09 (Miró-Pina et al., 2022). Interestingly, however, H3K9me3 and H3K27me3 could still be detected in the new MAC upon depletion of Gtsf1 (Figure 5A). Quantification of H3K9me3 and H3K27me3 fluorescence in the new developing MACs indicated that H3K9me3 and H3K27me3 accumulation is in fact significantly increased upon Gtsf1 depletion (Figure 5A, Figure S4). Additionally, we found that a functional tagged Ezl1, the catalytic subunit of PRC2 (Frapporti et al., 2019), accumulates in the maternal MAC, where it is normally barely detected, upon Gtsf1 depletion (Figure S4).

**Figure 5.**
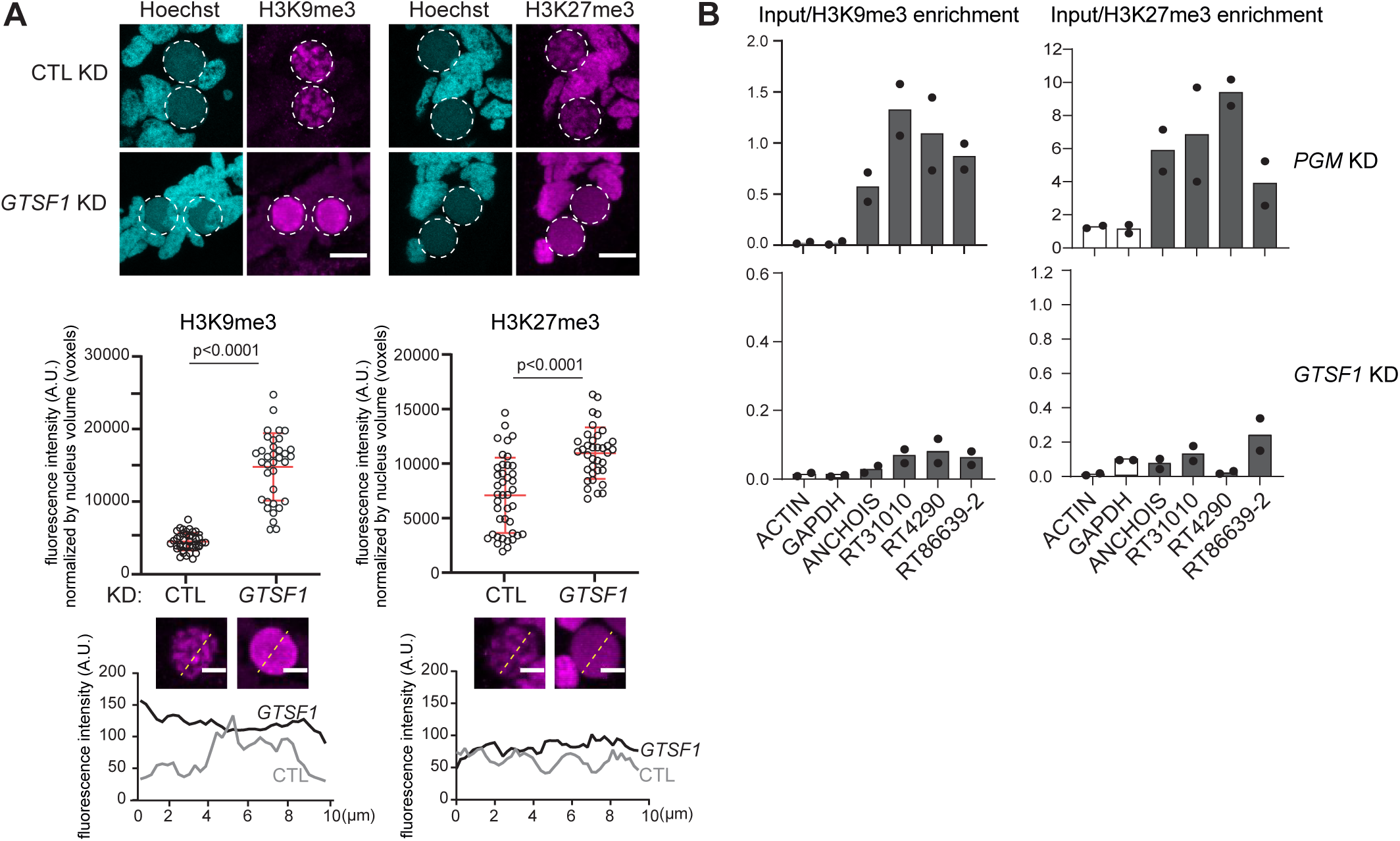
Gtsf1 depletion affects H3K9me3 and H3K27me3 levels and localization. A. H3K9me3 or H3K27me3 antibodies are used for immunostaining of wild type cells at T=15 hours after the onset of autogamy in *ICL7* (CTL) or *GTSF1* KD. Dashed white circles indicate the new developing MACs. The other Hoechst-stained nuclei are the fragments of maternal MAC and the MICs. Note that H3K9me3 staining is visible in the fragments of the maternal MAC upon *GTSF1* KD, while it is not in *CTL* KD. Scale bar, 10 μm. Middle: quantification of H3K9me3 and H3K27me3 fluorescence signal in the new developing MACs (see Materials and Methods). Number of nuclei > 30 in each condition. Bars correspond to mean ± SD. Mann-Whitney statistical tests. Bottom: quantification of H3K9me3 and H3K27me3 fluorescence signal along the line crossing the nucleus (yellow). B. Barplots of H3K9me3 and H3K27me3 enrichment over input (log2) for genes (*ACTIN*, *GAPDH*) (white) and TE copies (*ANCHOIS*, *RT31010*, *RT4290*, *RT86639-2*) (grey) determined by ChIP-qPCR upon *PGM* (control-2 replicates) and *GTSF1* KD (2 replicates).

In control conditions, H3K9me3 and H3K27me3 signals in new developing MACs display a diffuse pattern that gradually forms nuclear foci, as previously reported (Figures 5A) (Lhuillier-Akakpo et al., 2014). In Gtsf1-depleted cells, however, the H3K9me3 and H3K27me3 signal remains diffuse as development proceeds and foci are not detected (Figure 5A). Thus, despite the fact it is exclusively localized in the maternal MAC, Gtsf1 controls both the levels of Ezl1-mediated H3K9me3 and H3K27me3 and the clustering of these marks into nuclear foci in the new developing MACs, a function reminiscent of that reported for downstream effectors of the DNA elimination pathway (de Vanssay et al., 2020; Lhuillier-Akakpo et al., 2014).

To determine whether H3K9me3 and H3K27me3 are correctly targeted to TEs, we performed chromatin immunoprecipitation (ChIP) experiments (T = 50 hours) upon *GTSF1* KD, and upon *PGM* KD as a positive control. ChIP-qPCR analysis showed that TEs are enriched for both H3K27me3 and H3K9me3 in *PGM* KD conditions, while genes are not enriched for these marks (Figure 5B), as previously reported (Frapporti et al., 2019). By contrast, H3K9me3 and H3K27me3 enrichment were reduced for the TEs we analyzed upon *GTSF1* KD (Figure 5B). Thus, though Gtsf1 is confined to the maternal MAC, it appears to be required for proper targeting of H3K9me3 and H3K27me3 to TEs in the new developing MACs.

### Gtsf1 is required for scnRNA selection

Our data show that Gtsf1 affects H3K9me3 and H3K27me3 targeting and DNA elimination in the new developing MAC, yet it is exclusively present in the maternal MAC. This suggests that Gtsf1 plays a role in the maternal MAC during scnRNA selection. To investigate this possibility, we first evaluated steady-state scnRNA levels after depletion of Gtsf1 compared with control knockdown at 8 different time points during autogamy (Figure S1). This indicated that loss of Gtsf1 leads to increased total accumulation of scnRNAs (Figure 6A). In contrast, loss of Ptiwi01/09 leads to destablization of scnRNAs (Bouhouche et al., 2011).

**Figure 6.**
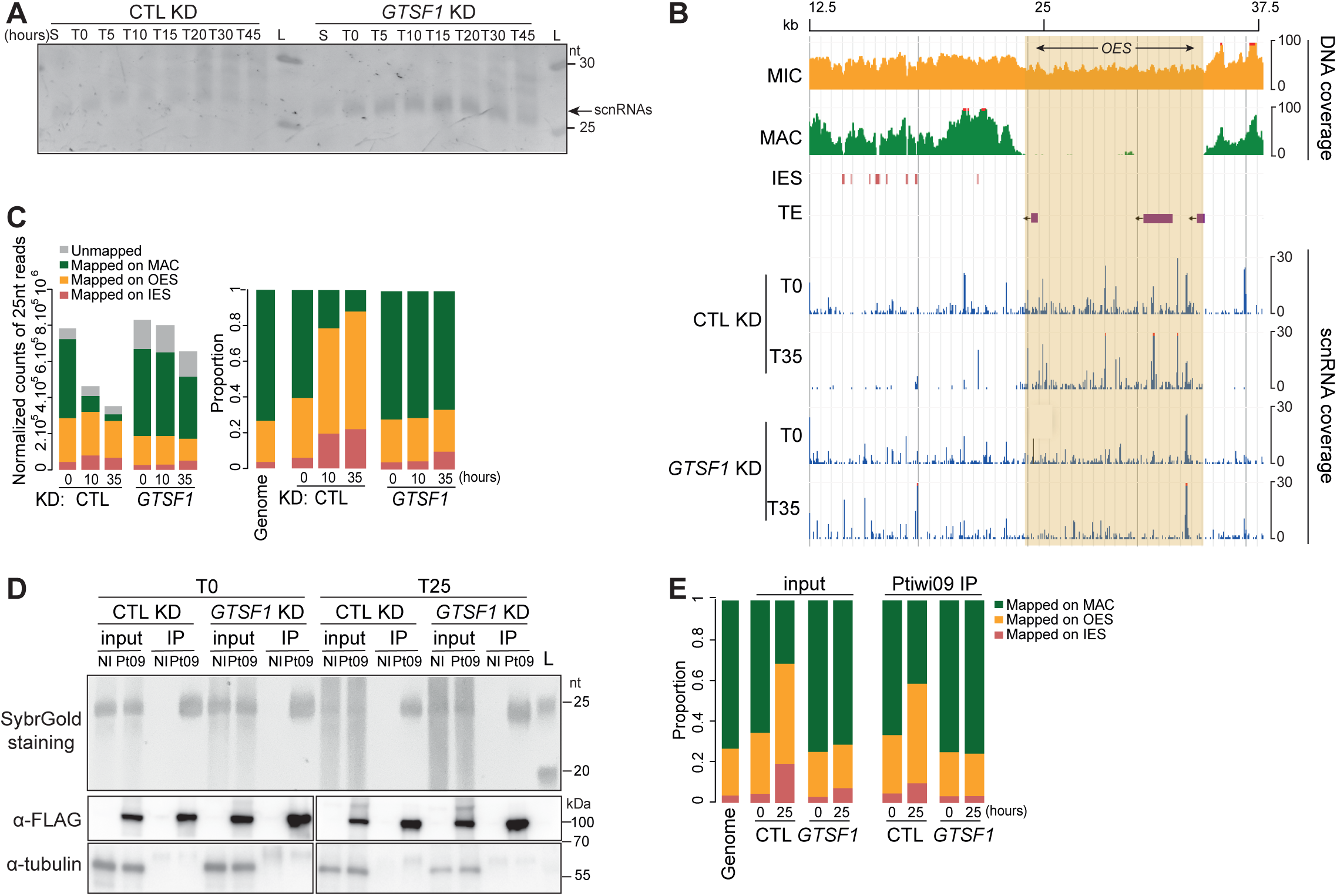
Gtsf1 is required for scnRNA selection. A. Gel electrophoresis of sRNAs from *ND7* (CTL) and *GTSF1* KD cells. Total RNA samples corresponding to different time points (S = starvation, T=0, 5, 10, 20, 30 and 45 hours after the onset of autogamy) were run on a denaturing 15% polyacrylamide-urea gel. After electrophoresis, the gel was stained with SybrGold. L: DNA low molecular weight marker (USB). The 25-nt scnRNAs are indicated. B. Representative genomic region depicting 25-nt scnRNA normalized coverage in *ICL7* (CTL) and *GTSF1* KD (NODE_1273_len-gth_39901_cov_28.471216 between 12.5 and 37.5 kb). C. Analysis of 25-nt scnRNA populations in *ICL7* (CTL) and *GTSF1* KD at different times points (T=0, 10 and 35 hours after the onset of autogamy). Bar plots show the normalized counts of 25-nt reads for each sample that map the MAC genome, IESs, or MIC-limited sequences (left) and the proportion of 25-nt reads for each category (right). Genome: proportion of each category (MAC, OES, IES) in the MIC genome. D. Upper panel: Gel electrophoresis of sRNAs before (input) or after (Ptiwi09 IP) immunoprecipitation of cells expressing *3X-FLAG-PTIWI09* and of non-injected cells (NI) at two different time points (T=0 and 25 hours after the onset of autogamy) in *ICL7* (CTL) and *GTSF1* KD. Total RNA samples were run on a denaturing 15% polyacrylamide-urea gel. After electrophoresis, the gel was stained with SYBR Gold. L: molecular weight marker (GeneRuler Ultra Low Range DNA ladder). Middle and lower panels: Western blot analysis of whole cell extracts from the same samples as the upper panel. Ptiwi09 detection was performed using FLAG antibodies and tubulin antibodies were used for normalization. E. Sequencing analysis of scnRNA populations before (input) or after (Ptiwi09 IP) immunoprecipitation at T=0 and 25 hours after the onset of autogamy in *ICL7* (CTL) or *GTSF1* KD (same samples as in panel D). Genome: proportion of each category (MAC, OES, IES) in the MIC genome.

We extended the scnRNA analysis by sequencing total small RNAs in the 20- to 30-nt range from Gtsf1-depleted cells at several time points (from T=0 to T=35 hours after the onset of autogamy) (Figure S1; Figure S5). In Gtsf1-depleted cells, the 25-nt scnRNAs are produced from the whole MIC genome at the beginning of the sexual cycle (T=0 hours), as in control RNAi conditions, consistent with previous work (Miró-Pina et al., 2023) (Figure 6B-C). This can be clearly observed when examining the mapping of 25-nt scnRNAs across an individual genomic region (Figure 6B), where MAC sequences, as well as MIC sequences (IES and OES), are covered. We therefore conclude that Gtsf1 is not involved in the process of scnRNA biogenesis. As autogamy proceeds (T=10 and T=35 hours), the proportion of 25-nt scnRNAs mapping to OES or IES over scnRNAs mapping to MAC sequences increases in control conditions (Figure 6B-C). At T=10 hours, scnRNAs mostly corresponded to MIC sequences (Figure 6C). However, MIC-specific scnRNAs (OES + IES) are not enriched over MAC-specific scnRNAs in Gtsf1-depleted cells, indicating an absence of MIC-specific scnRNA selection (Figure 6B-C; Figure S5). As illustrated for an individual genomic region, scnRNAs still cover IESs and the entire OES region, which comprises annotated transposable elements, at T=35 hours (Figure 6B). These patterns are in stark contrast to those observed upon loss of Ptiwi01/09 (Figure S5) and instead resemble those observed upon loss of PRC2-Ezl1 components (Miró-Pina et al., 2023; Wang et al., 2022). Importantly, because we still detect maternal MAC ncRNA production in the *GTSF1* knockdown experiments, the lack of scnRNA selection cannot be explained by a lack of maternal MAC ncRNA transcription (Figure S6).

To determine whether the scnRNAs corresponding to the entire germline genome that accumulate in Gtsf1*-*depleted cells are still loaded onto Ptiwi09, we performed Ptiwi09 IP followed by small RNA isolation (Furrer et al., 2017)(see Methods) upon *GTSF1* and control KD at two different time points (T=0 and T =25 hours) (Figure 6D; Figure S1). Sequencing of small RNAs revealed the presence of 25-nt scnRNAs in control, as expected (Furrer et al., 2017), and in Gtsf1-depleted cells at both time points (Figure S5). While the proportion of MIC-scnRNAs increases in the control at T=35 hours as expected, this is not the case upon Gtsf1 depletion, as the same proportions of MIC-and MAC-scnRNAs are detected in the IP at both time points, mirroring what is seen in the input (Figure 6E; Figure S5). This indicates that the non-selected scnRNAs that accumulate in Gtsf1-depleted cells are bound to Ptiwi09.

### Gtsf1 controls Ptiwi09 protein levels and ubiquitylation

To determine whether impairment of scnRNA selection could be explained by defects in the localization of scnRNAs, we analyzed the localization of the scnRNA-binding protein Ptiwi09 with a FLAG*-*tagged *PTIWI09* transgene (Miró-Pina et al., 2022). In control cells, the Ptiwi09 fusion protein is detected in the maternal MAC during MIC meiosis, then localized exclusively in the developing MAC (Figure 7A), as previously reported (Bouhouche et al., 2011). In cells lacking Gtsf1, the subcellular localization of Ptiwi09 proteins was unaffected (Figure 7A). Thus, the lack of MIC-specific scnRNA selection in the absence of Gtsf1 cannot be explained by the mis-localization of Ptiwi09-bound scnRNAs.

**Figure 7.**
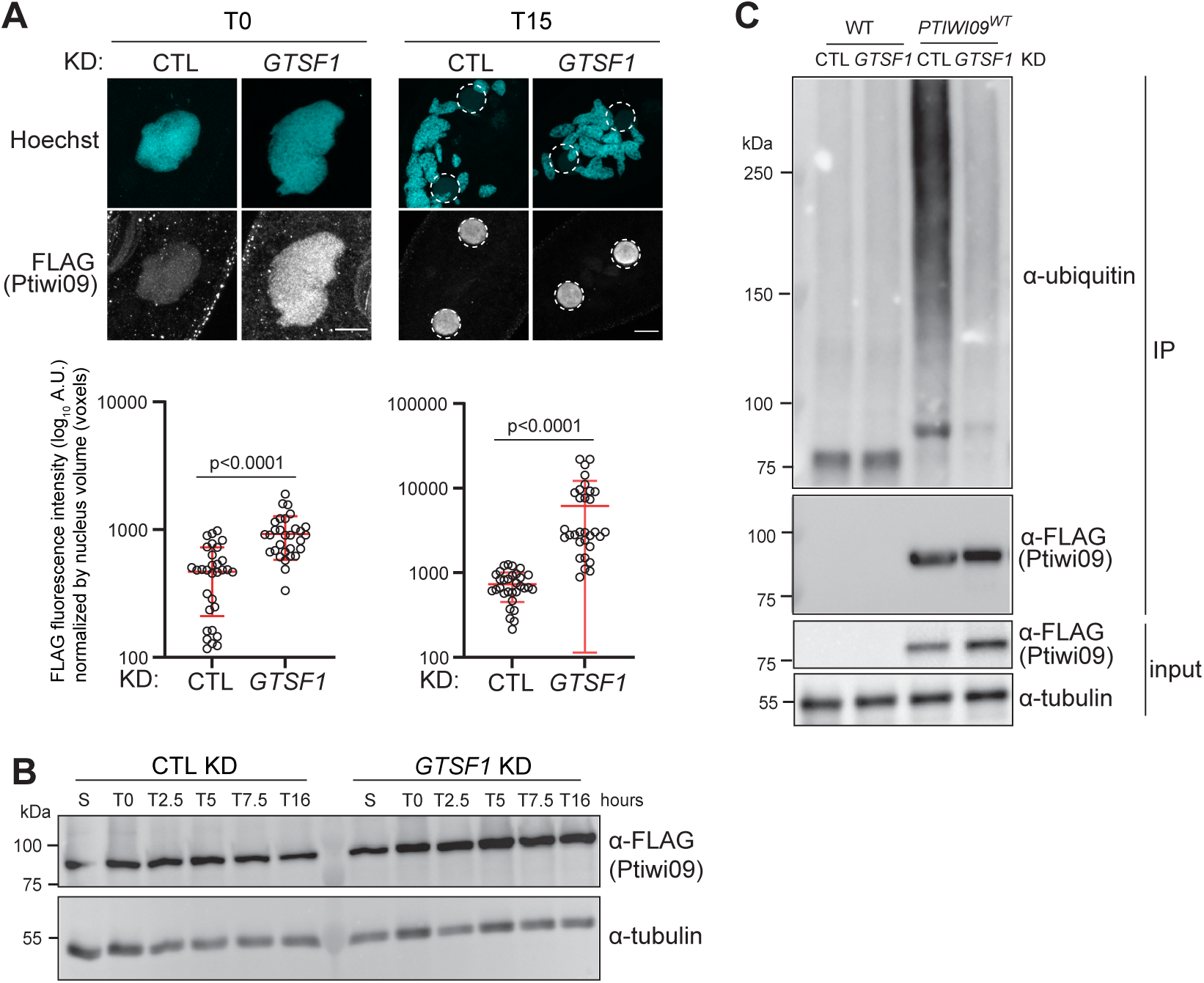
Gtsf1 controls Ptiwi09 protein levels and ubiquitylation. A. Anti-FLAG immunostaining of cells transformed with a *3xFLAG-PTIWI09* transgene at T=0 and T=15 hours after the onset of autogamy in *ICL7* (CTL) or *GTSF1* KDs. Dashed white circles indicates the new developing MACs. The other Hoechst-stained nuclei are the fragments of maternal MAC. Scale bar, 10 μm. Quantification of FLAG fluorescence signal in the nuclei (see Materials and Methods). Number of nuclei > 30 in each condition. Bars correspond to mean ± SD. Mann-Whitney statistical tests. B. Western blot analysis of whole cell extracts at different time points (S= starved; T=0; 2.5; 5; 7.5; 16 hours after the onset of autogamy) in *ND7* (CTL), *GTSF1* and *PTIWI01/09* KDs with FLAG antibodies to detect 3XFLAG-Ptiwi09 and tubulin antibodies for normalization. C. Western blot analysis of ubiquitylation of Ptiwi09 immunoprecipitation (IP) on wildtype cells (WT) and cells expressing *3XFLAG-PTIWI09* transgene (*PTIWI09^WT^*) at T=0 hours after the onset of autogamy upon *ICL7* (CTL) and *GTSF1* KD. Ptiwi09 detection was performed using FLAG antibodies before (input) and after (IP) Ptiwi09 immunoprecipitation. α-tubulin antibodies were used for normalization.

However, the intensity of the Flag signal was increased in Gtsf1-depleted cells. Therefore, we quantified the fluorescence intensity (see Materials and Methods) and found a significant increase of nuclear Ptiwi09 levels in the maternal MAC compared to control conditions (Figure 7A). Western bot analysis from whole cell extracts confirmed that the steady-state levels of the Ptiwi09 protein increase in Gtsf1-depleted cells (Figure 7B). Because the steady-state mRNA levels of *PTIWI01/09* are not impacted by *GTSF1* KD (Figure S2), we conclude that Gtsf1 controls Ptiwi09 protein accumulation more directly.

Given that the levels of the Ptiwi09 protein and the associated MAC-scnRNAs are increased in Gtsf1-depleted cells, we propose that Gtsf1 mediates the degradation of Ptiwi09/MAC-scnRNA complexes in the maternal MAC. This would explain the lack of MIC-scnRNA selection in the absence of Gtsf1. Given the general role of ubiquitin in triggering protein degradation (Varshavsky, 2017) and the reported case of ubiquitylation-mediated degradation of Argonaute protein (Kobayashi et al., 2018), we hypothesize that the ubiquitin pathway is involved in the degradation of Ptiwi09/MAC-scnRNA complexes. To test this possibility, we examined the levels of ubiquitylated proteins in Ptiwi09 IPs upon *GTSF1* and control KD, from cells collected at a stage when Ptiwi09 is detected in the maternal MAC (Figure 7C; Figure S1). Anti-ubiquitin antibodies revealed that Ptiwi09 and/or its associated proteins are ubiquitylated in a Gtsf1-dependent manner. Altogether, we conclude that Gtsf1 controls Ptiwi09 protein levels and the ubiquitylation of Ptiwi09 and/or its partners during scnRNA selection.

## Discussion

The nuclear *Paramecium* Gtsf1 homolog is an essential factor acting downstream of scnRNA biogenesis that is required for the selective degradation of a subpopulation of scnRNAs corresponding to MAC-destined sequences. As a result, scnRNA selection is defective in Gtsf1-depleted cells (Figure 6). Given its exclusive localization in the maternal MAC (Figure 3), the role of Gtsf1 in scnRNA selection strongly supports the idea that the maternal MAC is where selective degradation of scnRNAs occurs.

Like its counterparts in other organisms (Chen et al., 2014; Dönertas et al., 2013; Muerdter et al., 2013; Ohtani et al., 2013; Yoshimura et al., 2018, 2009), *Paramecium* Gtsf1 is required for Piwi-guided transcriptional silencing and repressive histone modifications at TE loci. In Gtsf1-depleted cells, TEs and a subset of IESs, which are enriched for long, scnRNA-and Ezl1-dependent sequences, are no longer eliminated in the new MAC (Figure 4; Figure S2). Some TEs are transcriptionally upregulated upon *GTSF1* KD, as previously reported for PRC2-Ezl1 components and *PTIWI01/09* KDs (Miró-Pina et al., 2022). Gtsf1 is thus a critical member of the DNA elimination pathway.

Other proteins involved in DNA elimination (Pdsg1, Ema1, Nowa1, PRC2 core complex and cofactors) are also necessary for scnRNA selection (Arambasic et al., 2014; Aronica et al., 2008; Miró-Pina et al., 2023; Swart et al., 2017; Wang et al., 2022). However, in contrast to Gtsf1, these proteins localize both in the maternal MAC and in the new MAC. Given their dual roles and localization, it is difficult to unambiguously assign them a specific function in each nucleus. We previously reported that the methyltransferase activity of PRC2-Ezl1 is essential for DNA elimination (Frapporti et al., 2019), while its function in scnRNA selection does not involve its catalytic activity (Miró-Pina et al., 2023). Gtsf1 is the only reported case described so far of a protein that is exclusively present in the maternal MAC and yet has an effect on DNA elimination events that take place in the new MAC. Thus, our analysis of Gtsf1 provides a unique demonstration that defective scnRNA selection ultimately impairs DNA elimination.

When scnRNA selection occurs normally, only Ptiwi09/MIC-scnRNA complexes are present in the new MAC (Figure 6; Figure S6) (Furrer et al., 2017). In contrast, when scnRNA selection is defective in Gtsf1-depleted cells, we show that Ptiwi09 proteins present in the new MAC are loaded with both MAC-and MIC-scnRNAs (Figure 6; Figure S6). We also find that H3K9me3 and H3K27me3 accumulate in the new MAC (Figure 5; Figure S6). Our ChIP data indicate that the Ptiwi09-scnRNA complexes guide the deposition of these repressive histone modifications along the genome with no specific enrichment on TEs (Figure 5). Surprisingly, this does not appear to induce DNA elimination, suggesting that the mere presence of the repressive histone modifications is not sufficient to elicit the introduction of DNA double-strand breaks. We speculate that a threshold local concentration of histone modifications must be reached to trigger elimination of the modified chromatin. Another possibility is that the lack of differentially methylated regions, and thus of boundaries between eliminated regions and flanking regions, precludes the identification of the correct sites for introduction of DNA breaks by the elimination machinery.

Despite the fact that GTSF1 is conserved, its functions appear to have diversified. In both *Drosophila* and mouse, GTSF1 is also necessary for piRNA-dependent TE silencing; however, the mechanisms appear to be quite different. In *Drosophila*, DmGTSF1 localizes to the nucleus, and is required for transcriptional transposon silencing but not for piRNA biogenesis (Dönertas et al., 2013; Muerdter et al., 2013; Ohtani et al., 2013). In mouse, by contrast, GTSF1 localizes to both the cytoplasm and the nucleus, and is essential for secondary albeit not primary piRNA production (Yoshimura et al., 2018). Mammalian GTSF1 appears to act as an auxiliary factor that potentiates the piRNA-directed RNA cleavage activities of PIWI proteins, transforming them into efficient endoribonucleases (Arif et al., 2022). In *C*. *elegans*, GTSF1 does not participate in the piRNA pathway, but is instead involved in the assembly of a complex that produces a specific class of endogenous small RNAs targeting genic transcripts (Almeida et al., 2018). The function we have uncovered here for Gtsf1 in *Paramecium,* where it controls the selective degradation of scnRNAs, represents yet another example that underscores the divergent contributions of GTSF1 proteins to small RNA silencing pathways. The somatic macronucleus, where Gtsf1 is exclusively localized in *Paramecium*, is devoid of TEs. Thus, the role of Gtsf1 in the control of TEs appears indirect. Like other Gtsf1 proteins, *Paramecium* Gtsf1 physically interacts with PIWI - the scnRNA binding protein Ptiwi09 (Figures 1-2) – but it has no effect on its localization (Figure 7; Figure S6), suggesting that Gtsf1 acts downstream of Ptiwi09. We showed that Gtsf1 controls the steady-state levels of the Ptiwi09 proteins and of its bound MAC-scnRNAs, which both accumulate in the nucleus upon *GTSF1* KD (Figure 7). We propose that Gtsf1 interacts with the particular sub-population of Ptiwi09 proteins that is loaded with MAC-scnRNAs and mediates the coordinated degradation of this Ptiwi09 subset as well as its bound MAC-scnRNA (Figure 8).

**Figure 8.**
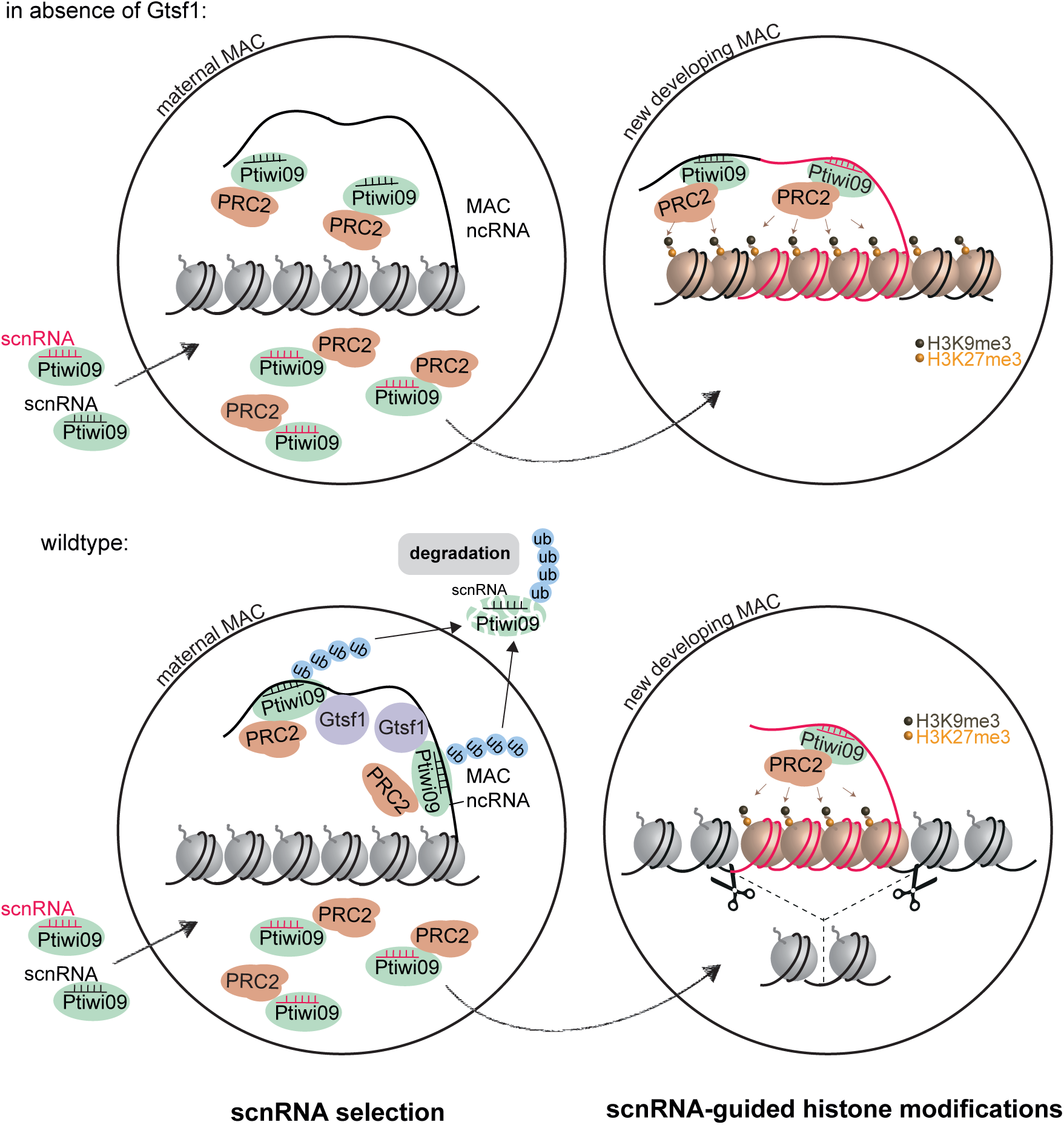
Model for the role of Gtsf1 in scnRNA selection. scnRNAs produced from the entire MIC genome during meiosis are bound to the Ptiwi09 protein and transported to the maternal macronucleus (MAC), where scnRNA selection occurs. The selective degradation of scnRNAs corresponding to MAC-destined sequences (black) results in the specific selection of the subpopulation corresponding to MIC-specific scnRNAs (in red). We propose that the Gtsf1 protein associates with the Ptiwi09 protein specifically when bound to scnRNAs (black) which are engaged with target nascent non-coding RNAs produced by the MAC genome (lower pannel). The physical interaction between Gtsf1, Ptiwi09-scnRNA complexes and non-coding MAC transcripts would trigger ubiquitylation of Ptiwi09, leading to degradation of Ptiwi09 and of its cognate scnRNAs (black). scnRNAs corresponding to MIC-specific sequences (red), which by definition cannot pair with MAC transcripts, are retained and trigger DNA elimination in the new developing MAC.

Indeed, our data are consistent with a scenario in which Gtsf1 is able to discriminate the pool of Ptiwi09 proteins that is bound to MAC-scnRNAs from the pool that is bound to MIC-scnRNAs. According to our model (Figure 8), Ptiwi09/MAC-scnRNA complexes are engaged in pairing interactions with non-coding RNA transcribed from the somatic maternal MAC genome, while the Ptiwi09/MIC-scnRNA complexes are not, because of the lack of sequence complementarity between MIC-scnRNAs and MAC transcripts. We envision that, as suggested in *M. musculus* and *Drosophila* (Ipsaro and Joshua-Tor, 2022), *Paramecium* Gtsf1 interacts with nascent transcripts from the maternal MAC to reinforce the association between Ptiwi09/MAC-scnRNA complexes and the targets. Importantly, however, while Gtsf1 may recognize Ptiwi09/scnRNA complexes that are already bound to their targets, it is also entirely possible that Gtsf1 could initiate target recognition and recruit Ptiwi09/MAC-scnRNA complexes rather than the converse.

How Ptiwi09 and its bound MAC-scnRNAs are degraded remains to be determined. Our data provide some hints that Gtsf1-mediated ubiquitylation might be critical to this process. Indeed, we found that Ptiwi09 and/or its associated proteins are ubiquitylated in a Gtsf1-dependent manner. Further work will be needed to identify the proteins whose ubiquitylation depends on Gtsf1. One possible mechanism is that the pairing of Ptiwi09/MAC-scnRNA complexes with nascent transcripts leads to a conformational change of the Ptiwi09 protein, allowing interaction with Gtsf1, which would then trigger Ptiwi09 ubiquitylation and degradation.

Target-directed degradation of scnRNAs during genome elimination in *Paramecium* exhibits striking similarities to recently reported instances of target-directed degradation of microRNAs in mammalian cells and *Drosophila* (Kingston et al., 2022; Wu and Zamore, 2021). miRNA degradation has been shown to rely on ubiquitylation and proteasomal degradation of the Argonaute protein (Han et al., 2020; Shi et al., 2020). Loss of scnRNA degradation upon Gtsf1 disruption is accompanied by excess accumulation of Ptiwi09, suggesting that an analogous mechanism may be at play during *Paramecium* genome elimination. These highly diverged phenomena of small-RNA removal may thus share basic commonalities. Future work aiming at unraveling the underlying mechanisms of target-directed scnRNA degradation might reveal that target-directed small RNA degradation is an ancient process that is more widespread than previously thought.

## Materials and Methods

### Paramecium strains, cultivation, and autogamy

All experiments were carried out with the entirely homozygous strain 51 of *P. tetraurelia*. Cells were grown in wheat grass powder (WGP) (Pines International) infusion medium bacterized the day before use with *Klebsiella pneumoniae*, unless otherwise stated, and supplemented with 0.8 mg/mL β-sitosterol (Merck). Cultivation and autogamy were carried out at 27°C as described (Beisson et al., 2010a, 2010b).

### Gene silencing experiments

Plasmids used for T7Pol-driven dsRNA production in silencing experiments were obtained by cloning PCR products from each gene using plasmid L4440 and *Escherichia coli* strain HT115 DE3, as previously described (Galvani and Sperling, 2002). Sequences used for silencing of ICL7a, GTSF1, EZL1, PTIWI01, PTIWI09, were segments 1-580 of PTET.51.1.G0700039 (ICL7a); 249-479 (GTSF1#1) or 32-441 (GTSF1#2) of PTET.51.1.G0490019 (GTSF1); 989-1501 of PTET.51.1.G1740049 (EZL1); 41-441 of PTET.51.1.G0710112 (PTIWI01); 50-439 of PTET.51.1.G0660118 (PTIWI09). Preparation of silencing medium and RNAi during autogamy were performed as described in (Baudry et al., 2009). Lethality of post-autogamous cells after RNAi was assessed by transferring 30–60 individual post-autogamous cells to standard growth medium. Cells with a functional new MAC were identified as normally growing survivors unable to undergo a novel round of autogamy if starved after ∼8 divisions. Cells usually divided two to four times before dying upon *GTSF1* KD as for *PTIWI01/09* KD, and unlike *EZL1* KD cells, which usually did not divide or only did so once before dying. See Figure S2 for details on RNAi-mediated KD experiments.

### Cytological stages monitoring and description

Progression of autogamy was followed by cytology with DAPI or Hoechst staining in the time course experiments. The progression through autogamy is not synchronous in the cell population (Berger, 1986). The time points refer to hours after T=0 hour (the onset of autogamy) that is defined as 50% of cells are autogamous (approximately 25% have a fragmented maternal MAC), as evaluated by cytological observation. See Figure S1 for details on progression of autogamy.

### Transformation with tagged transgenes

For the construction of in-frame *3xFLAG-HA-GTSF1* (pOC17)*, 3xFLAG-PTIWI09* (pJG091, Owsian et al., 2022) fusion plasmids, 3xFLAG-HA or 3xFLAG tags that were codon-optimized for the *P. tetraurelia* genetic code were added to the 5’ of the gene. As a result, the tag is fused to the N-terminus of *GTSF1* or of *PTIWI09*. The fusion proteins are expressed under the control of their endogenous regulatory regions (promoter and 3’UTR). *GTSF1* contains 25-bp upstream and 64-bp downstream of its open reading frame, and *PTIWI09* 257-bp upstream and 204-bp downstream. The *3xFLAG-HA-GTSF1* fusion transgene is RNAi-resistant. The 247-480 DNA fragments of *GTSF1* coding sequence was replaced by RF cloning with synthetic DNA sequences (Eurofins Genomics) designed to maximize nucleotide sequence divergence with the endogenous genomic loci without modifying the amino acid sequences of the encoded proteins. *3xFLAG-HA* (pIC12; 3xFLAG-HA tag only under *EZL1* regulatory sequences) *9/19/23 9:08:00 AM*(Miró-Pina et al., 2022), *3xFLAG-HA-EZL1* (pCM10) (Frapporti et al., 2019) and *3xFLAG-HA-GTSF1* (pOC17) were linearized by XmnI, *3xFLAG-HA-PTIWI09* (pAH30) (Miró-Pina et al., 2022) by SpeI, and *3xFLAG-PTIWI09* (pJG091) by AhdI for microinjection into the MAC of vegetative cells. No lethality was observed in the post-autogamous progeny of injected cells, indicating that none of the fusion constructs interfered with the normal progression of autogamy.

### Immunoprecipitation

Ptiwi09 immunoprecipitation (Figure 1; Table S1) was performed as follows: *Paramecium* cells transformed with the 3x*FLAG-PTIWI09* transgene as well as non-transformed cells were grown until ∼1.5 hours before T=0 hours. 200 mL of cells (4,000 cells/mL) were frozen in liquid nitrogen. The cell pellet (300 µL) was resuspended in 3 mL lysis buffer (50mM Tris pH 7.4, 300mM NaCl, 2mM MgCl2, 10% glycerol, 2mM EDTA, 0.3% Triton X-100, 2 mM PMSF, 1x Pierce Protease Inhibitor Tablets, EDTA-Free), kept on ice for 20 min then lysed with a Potter-Elvehjem homogenizer. To whole cell lysate is incubated with (100 U) TURBO™ DNase (Ambion) for 30 min at 4°C then centrifuged for 30 min at 20,817 g at 4°C. The supernatant was incubated with 40 μl anti-FLAG M2 agarose gel (A2220, Sigma) for 4 hours at 4°C. Beads were washed five times with lysis buffer and 5 times with wash buffer (10 mM Tris pH 7.4, 150 mM NaCl).

For the 3xFLAG-HA-Gtsf1 immunoprecipitation (Figure 2 and Table S1), *Paramecium* nuclear protein extracts were performed as previously described (Frapporti et al., 2019). 10^6^ autogamous cells (T=∼0 hours) were lysed with a Potter-Elvehjem homogenizer in 3 volumes of lysis buffer (10 mM Tris pH 6.8, 10 mM MgCl2, 0.2% Nonidet P-40, 1 mM PMSF, 4 mM benzamidine, 1x Complete EDTA-free Protease Inhibitor Cocktail tablets (Roche)). The nuclei-containing pellet was collected by centrifugation and washed with the addition of 2.5 volumes of washing solution (0.25M sucrose, 10 mM MgCl2, 10 mM Tris pH 7.4, 1 mM PMSF, 4 mM benzamidine, 1x Complete EDTA-free Protease Inhibitor Cocktail tablets (Roche)). The pellet was incubated in 1 volume of nuclear extraction buffer 2 × (100 mM Hepes pH 7.8, 100 mM KCl, 30 mM NaCl, 0.2 mM EDTA, 20% Glycerol, 2 mM DTT, 0.02% Nonidet P-40, 2 mM PMSF, 2x Complete EDTA-free Protease Inhibitor Cocktail tablets (Roche)) for 1 hour at 4°C. The salt-extractable fraction at 15 mM NaCl was recovered following centrifugation for 3 minutes at 10,000 g at 4°C. Nuclear extracts were incubated overnight at 4°C with 150 µl anti-FLAG M2 magnetic beads (M8823, Sigma) that were pre-washed with 1 mL TEGN buffer (20 mM Tris pH 8, 0.1 mM EDTA, 10% Glycerol, 150 mM NaCl, 0.01% Nonidet P-40). Beads were washed five times with TEGN buffer and eluted with 3xFLAG peptide (F4799, Sigma-Aldrich) (45 µl) and the same volume of TEGN buffer at 4°C for 5 hours.

### FLAG-Ptiwi09 immunoprecipitation for ubiquitylation analysis

200 mL (3,600 cells/mL) of *Paramecium* cells transformed with the 3x*FLAG-PTIWI09* transgene (Figure 7, Table S1) at T=0 hours were lysed in 3 mL cold lysis buffer (50 mM Tris pH 8, 300 mM NaCl, 1% Triton X-100, 0.01% NP-40, 10% glycerol, 5 mM EDTA, 25 mM N-Ethylmaleimide (Sigma), 1x Inhibitor cocktail cOmplete™ ULTRA Tablets EDTA free (Roche), 2mM PMSF) using a Potter-Elvehjem homogenizer until all nuclei were destroyed. The lysate was incubated for 1 hour at 4°C and centrifuged for 30 min at 18,400 g. The supernatant was incubated overnight at 4°C with 50 μl of anti-FLAG M2 magnetic beads (M8823, Sigma). The beads were washed five times with freshly prepared lysis buffer, two times 10 min with high salt buffer (10 mM Tris pH 7.4, 1 M NaCl), two times 10 min with 10mM Tris pH 7.4, 1M urea and finally two times for 10 min in 10 mM Tris pH 7.4, 150mM NaCl. The beads were boiled for 10 min at 95°C in 100 μl 1x LDS Sample Buffer (Invitrogen).

### Western blot and silver staining

For western blot, electrophoresis and blotting were carried out according to standard procedures. Samples were run on 10% Tris-Glycine Biorad gels. Blotting was performed overnight using a nitrocellulose (GE10600002, Merck) or PVDF membrane (Bio-Rad). FLAG (1:1,000 or 1:4,000) (MAI-91878, Thermo Fisher Scientific or F1804, Sigma), H3 (1:10,000) (AS- 61704, Eurogentec), α-tubulin (1:5,000) (sc-8035, Santa Cruz) or TEU435 (1:4,000)(Callen et al., 1994) were used for primary antibodies. Secondary horseradish peroxidase-conjugated anti-mouse or anti-rabbit IgG antibodies (Promega) were used at 1: 2,500 or 1: 8,000 dilution followed by detection by ECL.

To detect ubiquitin (Figure 7C), samples were run on NuPage 3-8% Tris-Acetate Gel at 4°C 90V. Blotting was performed overnight using a PVDF membrane (F1804, Sigma). The membrane was boiled for 30 min in water and incubated for 1 hour in 5% skim milk in TBS pH 7.4 then for 2 hours in primary antibody (1:2,000 ubiquitin monoclonal antibody 13-1600 Invitrogen) and for 1 hour at room temperature in secondary antibody (1:4,000 anti-Mouse-HRP, Promega). In order to detect the endogenous Ptiwi09 protein, custom antibodies were raised. The QLANTEIVNKKAGTK peptide of Ptiwi09 was used for rabbit immunization (Eurogentec). Polyclonal antibodies were purified by antigen affinity purification. Antibody α-Ptiwi09-433 specificity was tested on western blot (1:2,000) using whole cell extracts.

Silver staining was carried out with SilverQuest (Invitrogen LC6070) according to the manufacturer’s instructions.

### Mass spectrometry

#### Sample preparation

For Ptiwi09 immunoprecipitation, the agarose beads were analyzed at the Mass Spectrometry Laboratory at the Institute of Biochemistry and Biophysics PAS. At first, cysteines were reduced by 1 hour incubation with 20 mM Tris(2-carboxyethyl)phosphine (TCEP) at 60°C followed by 10 min incubation at room temperature with 50 mM methyl methanethiosulfonate (MMTS). Digestion was performed overnight at 37°C with 1 µg of trypsin (Promega). The tryptic digestion was stopped by lowering the pH of the reaction below pH 4 by adding extraction buffer (0.1% TFA 2% acetonitrile). The agarose beads were separated from solution by centrifugation. The resulting peptide mixtures were applied to RP-18 pre-column (Waters, Milford, MA) using water containing 0.1% FA as a mobile phase and then transferred to a nano-HPLC RP-18 column (internal diameter 75 µM, Waters, Milford MA) using ACN gradient (0 – 35% ACN in 160 min) in the presence of 0.1% FA at a flow rate of 250 nl/min. The column outlet was coupled directly to the ion source of an Orbitrap Elite mass spectrometer (Thermo Electron Corp., San Jose, CA) working in the regime of data-dependent MS to MS/MS switch. A blank run ensuring absence of cross-contamination from previous samples preceded each analysis.

For Gtsf1 immunoprecipitation, gel plugs were discolored using a solution of ACN/NH4HCO3 50 mM (50/50) for 15 minutes with agitation, reduced with a 10-mM DTT solution for 45 minutes at 56°C, then alkylated using a 55-mM IAA solution for 45 minutes at room temperature. After a washing and dehydration step, proteins in the plugs were digested overnight with trypsin (Promega) at 37°C in a 25-mM NH4HCO3 buffer (0.2 µg trypsin in 20 µL). The digested peptides were loaded and desalted on evotips provided by Evosep (Odense, Denmark) according to the manufacturer’s instructions before LC-MS/MS analysis at the Mass Spectrometry Facility at Institut Jacques Monod. The samples were analyzed on a timsTOF Pro 2 mass spectrometer (Bruker Daltonics, Bremen, Germany) coupled to an Evosep one system (Evosep, Odense, Denmark) operating with the 30SPD method developed by the manufacturer. Briefly, the method is based on a 44-min gradient and a total cycle time of 48 min with a C18 analytical column (0.15 x 150 mm, 1.9µm beads, ref EV-1106) equilibrated at 40°C and operated at a flow rate of 500 nL/min. H2O/0.1 % FA was used as solvent A and ACN/0.1 % FA as solvent B. The timsTOF Pro 2 was operated in PASEF mode1 over a 1.3 sec cycle time. Mass spectra for MS and MS/MS scans were recorded between 100 and 1700 m/z. Ion mobility was set to 0.75-1.25 V·s/cm2 over a ramp time of 180 ms. Data-dependent acquisition was performed using 6 PASEF MS/MS scans per cycle with a near 100% duty cycle. Low m/z and singly charged ions were excluded from PASEF precursor selection by applying a filter in the m/z and ion mobility space. The dynamic exclusion was activated and set to 0.8 min, a target value of 16000 was specified with an intensity threshold of 1000. Collisional energy was ramped stepwise as a function of ion mobility.

#### Data analysis

MS raw files were processed using PEAKS Online X (build 1.8, Bioinformatics Solutions Inc.). Data were searched against the ParameciumDB database (*P. tetraurelia* protein annotation v2.0, download 2021_10, total entries 40460). Parent mass tolerance was set to 10 ppm for Ptiwi09 IP and 25 ppm for Gtsf1 IP, the fragment mass tolerance to 0.05 Da. Specific tryptic cleavage was selected and a maximum of 2 missed cleavages was authorized. For identification, the following post-translational modifications were included: oxidation (M) and deamidation (NQ) as variables and beta-methylthiolation (C) as fixed. Identifications were filtered based on a 1% FDR (False Discovery Rate) threshold at PSM level. Label free quantification was performed using the PEAKS Online X quantification module, allowing a mass tolerance of 20 ppm and a retention time shift tolerance of 1 min for match between runs for Ptiwi09 IPs and a CCS error tolerance of 0.05 and a retention time shift tolerance in autodetect for match between runs for Gtsf1 IP. Protein abundance was inferred using the top N peptide method and TIC was used for normalization. Multivariate statistics on proteins were performed using Qlucore Omics Explorer 3.8 (Qlucore AB, Lund, SWEDEN). A positive threshold value of 1 was specified to enable a log2 transformation of abundance data for normalization i.e. all abundance data values below the threshold will be replaced by 1 before transformation. The transformed data were finally used for statistical analysis i.e. evaluation of differentially present proteins between two groups using a Student’s bilateral t-test and assuming equal variance between groups. A p-value better than 0.05 was used to filter differential candidates. The mass spectrometry proteomics data have been deposited to the ProteomeXchange Consortium via the PRIDE (Perez-Riverol et al., 2022) partner repository with the dataset identifiers PXD045266 (Ptiwi09) and PXD045214 (Gtsf1).

#### Immunofluorescence and quantification

As described in (Frapporti et al., 2019), cells were fixed for 30 min in solution I (10 mM EGTA, 25 mM HEPES, 2 mM MgCl2, 60 mM PIPES pH 6.9, PHEM 1X; 1% formaldehyde, 2.5% Triton X- 100, 4% sucrose), and for 10 min in solution II (PHEM 1X, 4% formaldehyde, 1.2% Triton X-100, 4% sucrose). Following blocking in 3% bovine serum albumin-supplemented Tris buffered saline-Tween 20 0.1 % (TBST) for 10 min, fixed cells were incubated overnight at room temperature under agitation with primary antibodies as follows: rabbit anti-H3K9me3 (1:200) (Frapporti et al., 2019), rabbit anti-H3K27me3 (1:1,000) (Frapporti et al., 2019), and mouse anti-FLAG (1:200) (MAI-91878, Thermo Fisher Scientific). Cells were labeled with Alexa Fluor 568-conjugated goat anti-rabbit IgG, Alexa Fluor 488-conjugated goat anti-rabbit IgG or Alexa Fluor 568-conjugated goat anti-mouse IgG at 1:500 for 1 hour, stained with 1 μg/mL Hoechst for 5–10 min and finally mounted in Citifluor AF2 glycerol solution. Images were acquired using a Zeiss LSM 780 or 980 laser-scanning confocal microscope and a Plan-Apochromat 63 × /1.40 oil DIC M27 objective. Z-series were performed with Z-steps of 0.35 μm. Quantification was performed as previously described (Vanssay et al., 2020) using ImageJ. The volume of the nucleus (in voxels) was estimated as follows: using the Hoechst channel, the top and bottom Z stacks of the developing MAC were defined to estimate nucleus height in pixels. The equatorial Z stack of the developing MAC was defined, and the corresponding developing MAC surface was measured in pixels. The estimated volume of the developing MAC was then calculated as the product of the obtained nucleus height by the median surface. For each Z stack of the developing MAC, the H3K9me3 or H3K27me3 fluorescence intensity was measured and corrected using the ImageJ "subtract background" tool. The sum of the corrected H3K9me3, H3K27me3 or FLAG fluorescence intensities for all the Z stacks, which corresponds to the total H3K9me3, H3K27me3 or FLAG fluorescence intensity, was divided by the estimated volume to obtain the H3K9me3, H3K27me3 or fluorescence intensity per voxel in each nucleus. For each condition at least 30 nuclei were quantified. Mann-Whitney statistical tests were performed with GraphPad Prism. All of the statistical details of experiments can be found in the figure legends.

#### Chromatin immunoprecipitation (ChIP)

ChIP experiments were performed with H3K9me3 or H3K27me3 antibodies as previously described (Frapporti et al., 2019). From ChIP-enriched samples and inputs, DNA was extracted with phenol, precipitated with glycogen in sodium acetate and ethanol and resuspended in deionized distilled water. Enrichment compared to input was analyzed by qPCR. qPCR was performed using the ABsolute QPCR SYBR Green Capillary Mix (Thermo Fisher Scientific) on the LightCycler 1.5 thermal cycling system (Roche). qPCR amplification was done with primers listed in Table S4.

#### DNA extraction and sequencing

DNA for deep-sequencing was isolated from post-autogamous cells (T=∼50 hours or T=60 hours) as previously described (Arnaiz et al., 2012). Briefly, cells were lysed with a Potter-Elvehjem homogenizer in lysis buffer (0.25 M sucrose, 10 mM MgCl2, 10 mM Tris pH 6.8, 0.2% Nonidet P-40). The nuclei-containing pellet was washed with washing buffer (0.25 M sucrose, 10 mM MgCl2, 10 mM Tris pH 7.4), loaded on top of a 3-mL sucrose layer (2.1 M sucrose, 10 mM MgCl2, 10 mM Tris pH 7.4) and centrifuged in a swinging rotor for 1 hr at 210,000 g. The nuclear pellet was collected and diluted in 200 μl of washing buffer prior to addition of three volumes of proteinase K buffer (0.5 M EDTA pH 9, 1% N-lauryl sarcosine sodium, 1% SDS, 1 mg/mL proteinase K). Following overnight incubation at 55°C, genomic DNA was purified and treated with RNase A.

DNA from sorted new MACs subjected to deep sequencing was obtained using fluorescence-activated nuclear sorting at T=25 hours after the onset of autogamy upon *GTSF1* or CTL KD, as described in (Zangarelli et al., 2022). Briefly, nuclei were collected from a 500 mL culture of autogamous cells at T=25 hours and subjected to flow cytometry sorting. 60,000 to 100,000 new MACs were sorted based upon their PgmL1 labelling (Zangarelli et al., 2022), and used for subsequent genomic extraction using QIAamp DNA micro kit (Qiagen). A microscope slide with 500 sorted nuclei was prepared in parallel to check for nuclear integrity and ploidy (Figure S3A) and anti-Pgm antibodies (Pgm 2659 GP, Dubois et al., 2017) were used in Figure S3B. Genomic DNA libraries were prepared either with the Westburg NGS DNA Library prep kit, or with the KAPA DNA HyperPrep (Kapa Biosciences, KK8504) and adapters IDT for Illumina TruSeq DNA UD Indexes (Illumina 20022370), according to the manufacturer recommendations. The quality of the final libraries was assessed with an Agilent Bioanalyzer, using an Agilent High Sensitivity DNA Kit. Library concentration was determined by qPCR using the Kapa Library Quantification kit (Kapa Biosciences, KK4824), according to manufacturer instructions. Libraries were pooled in equimolar proportions and sequenced using paired-end 2×75 pb runs, on an Illumina NextSeq500 instrument, using NextSeq 500 High Output 150 cycles kit for the samples, or using paired-end 2×100 bp runs on an Illumina NovaSeq 6000 instrument, using NovaSeq 6000 S1 Reagent Kit (200 cycles), (Illumina, 20028318) with the 0,5% addition of control library Phix (Illumina, FC-110-3001). Sequencing metrics are available in Table S3.

#### RNA extraction and sequencing

Total RNA samples were extracted as previously described (Frapporti et al., 2019) from 200– 400 mL of culture at 500 cells/mL for vegetative cells or at 2,000–4,000 cells/mL at different time-points during autogamy. Briefly, cells were centrifuged and flash-frozen in liquid nitrogen prior to TRIzol treatment, modified by the addition of glass beads for the initial lysis step. Alternatively, long and short RNA molecules were isolated from *Paramecium* cells using RNAzol RT (MRC). Total RNA quality was assessed with an Agilent Bioanalyzer 2100, using Agilent RNA 6000 pico kit. Directional polyA RNA-Seq libraries were constructed using the TruSeq Stranded mRNA library prep kit, following the manufacturer’s instructions. The quality of the final libraries was assessed with an Agilent Bioanalyzer, using a High Sensitivity DNA Kit. Libraries were pooled in equimolar proportions and sequenced using paired-end 2×75 pb runs, on an Illumina NextSeq500 instrument, using NextSeq 500 High Output 150 cycles kit.

Small RNAs of 15-35 nt were purified from total RNA on a 15% TBE/Urea gel. Alternatively, small RNAs were first enriched using RNAzol RT (MRC) from frozen cells then purified on gels. Small RNA libraries were constructed using the NEBNext Small RNA kit or TruSeq Small RNA Library Prep (Illumina, RS-200-0012, RS-200-0036) according to the manufacturer recommendations. The quality of the final libraries was assessed with an Agilent Bioanalyzer, using an Agilent High Sensitivity DNA Kit. Libraries were pooled in equimolar proportions and sequenced using a single read 75 bp run on an Illumina NextSeq500 instrument, using NextSeq 500 High Output 75 cycles kit, or paired-end 100 bp run on an Illumina NovaSeq 6000 instrument, using NovaSeq 6000 S1 Reagent Kit (Illumina, 20028318). Sequencing metrics are available in Table S3.

#### RNA Immunoprecipitation

200 mL of cells were collected at T=0 and T=25 hours after the onset of autogamy (Figure 6, Table S1) and RNA was isolated using RNazol RT. 5 µg of small RNA fraction was run on 15% TBE/Urea gel using Owl™ S4S sequencing gel system (Thermo Scientific) and small RNAs of 20- 35 nt were gel-purified for library construction (input). FLAG-Ptiwi09 immunoprecipitation was performed as described in (Furrer et al., 2017). After immunoprecipitation, RNA was extracted with phenol:chlorophorm, precipitated, dried, resuspended and denatured in Gel Loading Buffer II (AM8546G Thermo Fisher Scientific), and run on a 15% TBE/Urea gel. The gel was stained with SYBR Gold Nucleic Acid Gel Stain. Small RNAs of 20-35 nt were gel-purified and used for library construction (IP).

#### RT-PCR

5 µg of large RNA fraction isolated from frozen cells with RNAzol RT (MRC) was treated with TURBO® DNase (Thermo Fischer Scientific), extracted with phenol pH 4.3, then with chlorophorm, and precipitated. RNA concentration was estimated using Nanodrop (Thermo Fisher Scientific). 1 µg of RNA was reversed-transcribed using RevertAid H Minus Reverse Transcriptase (Thermo Fischer Scientific) and random hexamer. PCR amplification was done with primers listed in Table S4.

#### Analysis of sequencing data

Sequencing data were demultiplexed using CASAVA (v1.8.2) and bcl2fastq2 (v2.18.12), then adapters were removed using cutadapt (v3.4). Reads were mapped using Bowtie2 (v2.2.9) on known contaminants (mitochondrial genomes, ribosomal DNA, and bacterial genomes). The sequencing data were mapped on *P. tetraurelia* strain 51 MAC (ptetraurelia_mac_51.fa), MAC+IES (ptetraurelia_mac_51_with_ies.fa) and MIC (ptetraurelia_mic2.fa) reference genomes using Bowtie2 (v2.2.9 --local -X 500), Hisat2 (v2.1.0, --rna-strandness FR --min-intronlen 20 --max-intronlen100) or BWA (v0.7.15 -n 0) for DNA-seq, mRNA-seq or sRNA-seq data respectively. For genome browser screenshots, the sequencing coverage was normalized using deeptools (bamCoverage v3.2.1 --binSize 1 --normalizeUsing CPM). Gene annotation v2.0 (ptetraurelia_mac_51_annotation_v2.0.gff3), IES annotation v1 (internal_eliminated_sequence_PGM_ParTIES.pt_51.gff3) and TE annotation (ptetraurelia_mic2_TE_annotation_v1.0.gff3) were used in this study. All files are available from the ParameciumDB download section (https://paramecium.i2bc.paris-saclay.fr/download/Paramecium/tetraurelia/51/) (Arnaiz et al., 2020). R (v4.0.4) packages were used to generate images (ggplot2 v3.3.5; ComplexHeatmap v2.6.2; GenomicRanges v1.42; rtracklayer v1.50, seqinr v4.2-8, circlize v0.4.13, FactoMineR v2.4). Sequencing metrics are available in Table S3.

IES retention was evaluated using ParTIES (MIRET module v1.05 default parameters).

The RPKM coverage on TEs or genes was calculated using the reads counts, determined by htseq-count (v0.11.2, --mode=intersection-nonempty; for RNA-seq the option --stranded=yes has been used) on filtered BAM files (samtools v1.3.1 -q 30) then normalized by the number of mapped reads on the MIC genome.

The sRNA reads (20 to 30 nt) were successively mapped on the MAC, MAC+IES and MIC genomes to attribute them to a specific genome compartment: MAC-destined sequence (MAC), Internal Eliminated Sequences (IES) or Other Eliminated Sequences (OES). The 23nt siRNA reads that map to the RNAi targets were not considered. Normalized read counts were calculated using the number sequenced reads with a G+C content < 50%, compatible with a *Paramecium* G+C genomic content (∼27%).

## Funding

This work was supported by the Centre National de la Recherche Scientifique, the Agence Nationale pour la Recherche (ANR) [project “LaMarque” ANR-18-CE12-0005-02 to SD and MB]; [project “POLYCHROME” ANR-19-CE12-0015 to SD and OA]; [project “SELECTION” ANR-23- CE12-XXXX to SD and GC]; [project “CURE” ANR-21-CE12-0019-01 to MB]; the LABEX Who Am I? to SD (ANR-11-LABX-0071; ANR-11-IDEX-0005-02); the Fondation de la Recherche Médicale “Equipe FRM EQU202203014643” to SD; “FRM EQU202103012766” to MB; the National Science Centre of Poland [2019/32/C/NZ2/00472 to JG]; [2019/33/B/NZ2/01062 to JN]. OC was recipient of PhD fellowships from Université Paris Cité and Fondation de la Recherche Médicale (FDT202204014919) and of a EUR G.E.N.E. transition postdoc fellowship (ANR-17- EURE-0013; Université Paris Cité IdEx #ANR-18-IDEX-0001 funded by the French Government through its “Investments for the Future” program). JG, KN and JN received exchange grants from the CNRS and the Polish Academy of Sciences in the framework of the CNRS International Research Network (IRN) - Genome Editing in Ciliates (GEiC).

## Acknowledgements

We thank Daniel Holoch, Leticia Koch Lerner and Julien Richard Albert for critical reading of the manuscript and members of the Duharcourt lab for stimulating discussions. We thank Vinciane Régnier for advice for new MAC sorting. We acknowledge the ImagoSeine facility, member of the FranceBioImaging infrastructure supported by the ANR-10-INSB-04, and the Imagerie-Gif core facility, supported by the Agence Nationale de la Recherche (ANR-11-EQPX- 0029/Morphoscope, ANR-10-INBS-04/FranceBioImaging, ANR-11-IDEX-0003-02/ Saclay Plant Sciences). The present work has benefited from the sequencing of the I2BC High-throughput sequencing facility, supported by France Génomique (funded by the French National Program "Investissement d’Avenir" ANR-10-INBS-09) and the Genomics Core Facility CeNT UW, using NovaSeq 6000 platform financed by Polish Ministry of Science and Higher Education (decision no.6817/IA/SP/2018 of 2018-04-10), and the Mass Spectrometry Facilities at IBB PAN and at Institut Jacques Monod.

## Author contributions

OC and JG conducted most experiments; OA designed and performed the bioinformatic analyses of NGS data; CZ and MB performed new MAC sorting; KN performed ubiquitylation analysis; VL and GC analyzed the MS data; OC, JG, JN, SD designed the experiments. OC prepared most figures and SD wrote the paper with input from all authors.

## Notes

### Competing Interest Statement

The authors have declared no competing interest.

