## Supplementary Figures and Tables for "The nuclear PIWI-interacting protein Gtsf1 controls the selective degradation of small RNAs in *Paramecium*"

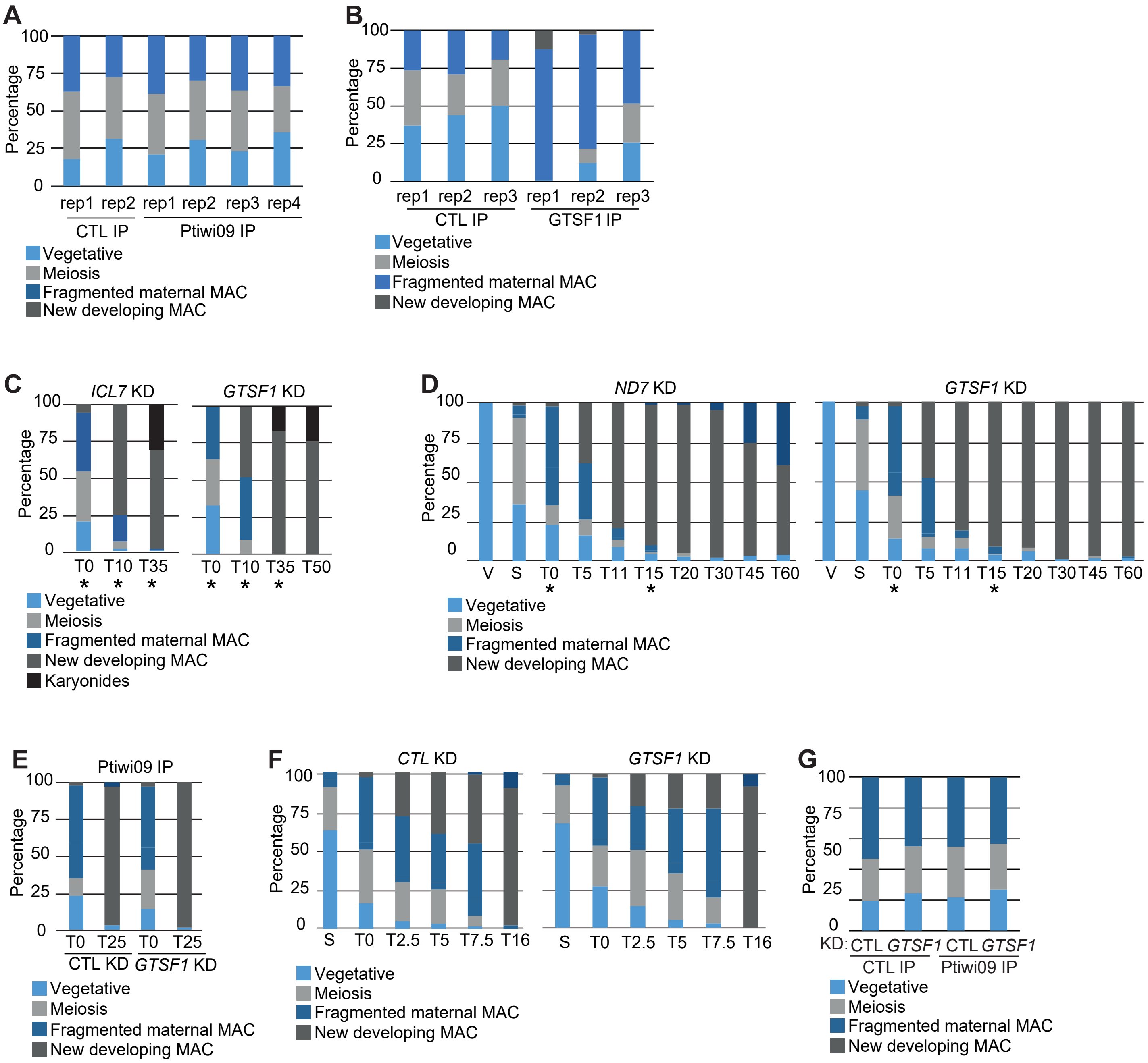

**Figure S1. Cytology of time course experiments**

Progression of autogamy followed by cytology with DNA staining in the time course experiments. >100 cells were counted in each condition.

A. Cytology of Ptiwi09 IP experiments, Related to Figure 1

B. Cytology of Gtsf1 IP experiments, Related to Figure 2

C. Cytology of time course experiment for sRNA and RNA sequencing in CTL (*ICL7* KD from Miró-Pina et al., 2022 and Miró-Pina et al., 2023) and *GTSF1* KD, related to Figures 4F, 6C, S2 and S5. The star (*) indicates the samples used for sRNA sequencing of replicate 1 (R1), related to Figure 6C and S5.

D. Cytology of time course experiment to detect scnRNA levels in CTL or *GTSF1* KD, related to Figure 6A and S5. The star (*) indicates the samples used for sRNA sequencing of replicate 2 (R2), related to Figure 6C and S5.

E. Cytology of time course experiment for sRNA sequencing from Ptiwi09 IP in CTL or *GTSF1* KD, related to Figure 6D, 6E and S5.

F. Cytology of time course experiment for Ptiwi09 detection in CTL or *GTSF1* KD, related to Figure 7B.

G. Cytology of time course experiment for Ptiwi09 IP in CTL or *GTSF1* KD, related to Figure 7C.

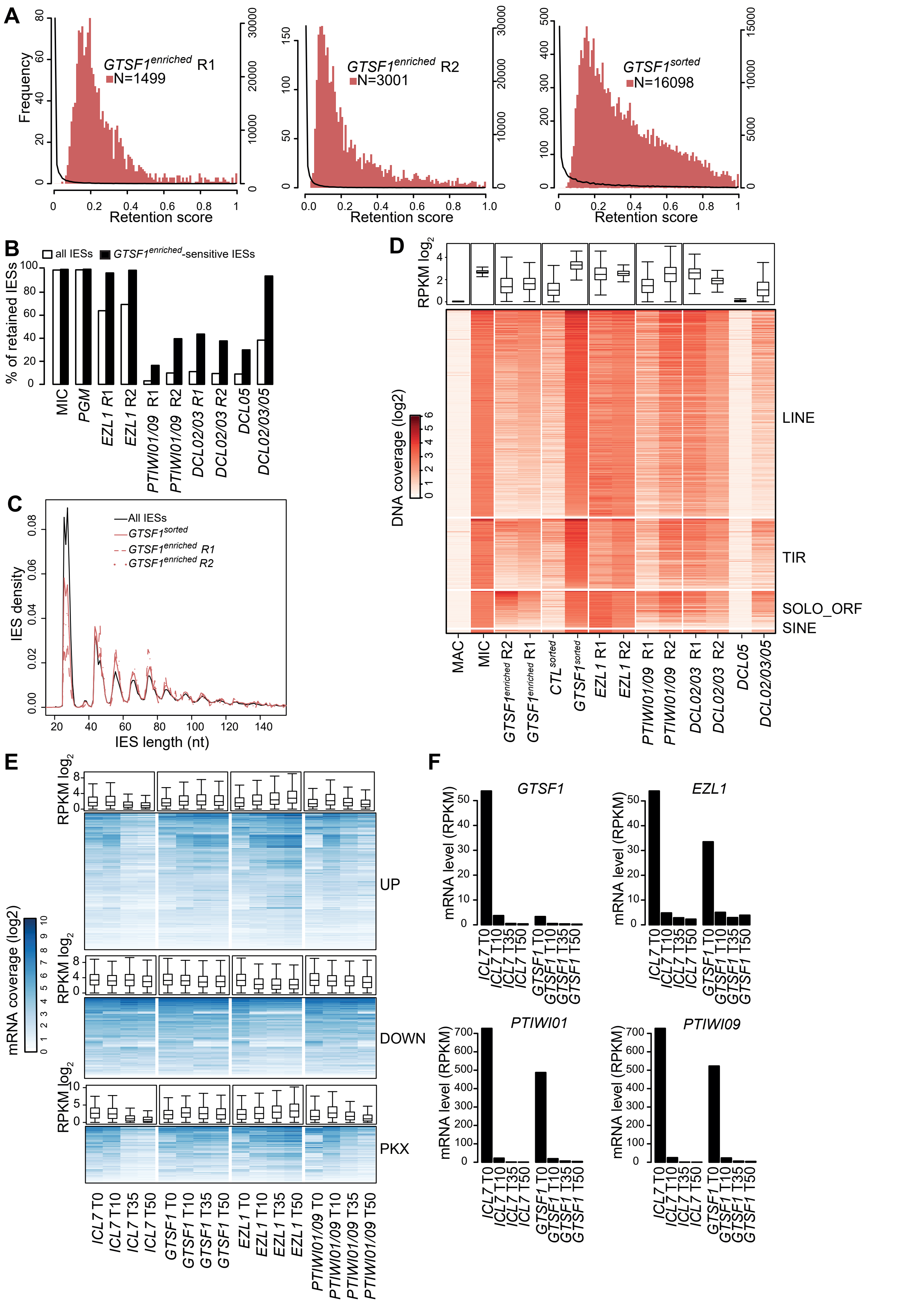

**Figure S2. Gtsf1 is required for efficient DNA elimination and TE silencing**

A. Histograms of IES retention scores for *GTSF1* KDs. The significantly retained IESs in *GTSF1* KD (2 replicates of *GTSF1*-enriched and 1 replicate of *GTSF1*-sorted) are represented by the red histograms (scale on the left), while the global distribution for all IESs retained in *GTSF1* KD is represented by the black curve (scale on the right).
B. Histogram of the percentage of retained IESs in MIC, *EZL1*, *PTIWI01/09*, *DCL02/03*, *DCL05* and *DCL02/03/05* KDs.
C. IES length distribution for all IESs and IESs retained upon *GTSF1* KD. Note that short IESs are under represented in *GTSF1* KD.
D. Heatmaps of TE normalized DNA coverage (Table S3). TE copies are ordered by the mean coverage of *GTSF1*-enriched R1, *GTSF1*-enriched R2 and *GTSF1*-sorted in each family.
E. Heatmaps of gene normalized RNA coverage for up- and down- regulated genes in *EZL1* KD (Frapporti et al., 2019) and up-regulated genes in *PGM-KU80C-XRCC4* KD (PKX) (Bazin-Gélis et al, 2023)
F. Barplots of *GTSF1*, *EZL1*, *PTIWI01* and *PTIWI09* mRNA levels in *ICL7* (CTL) and *GTSF1* KDs.

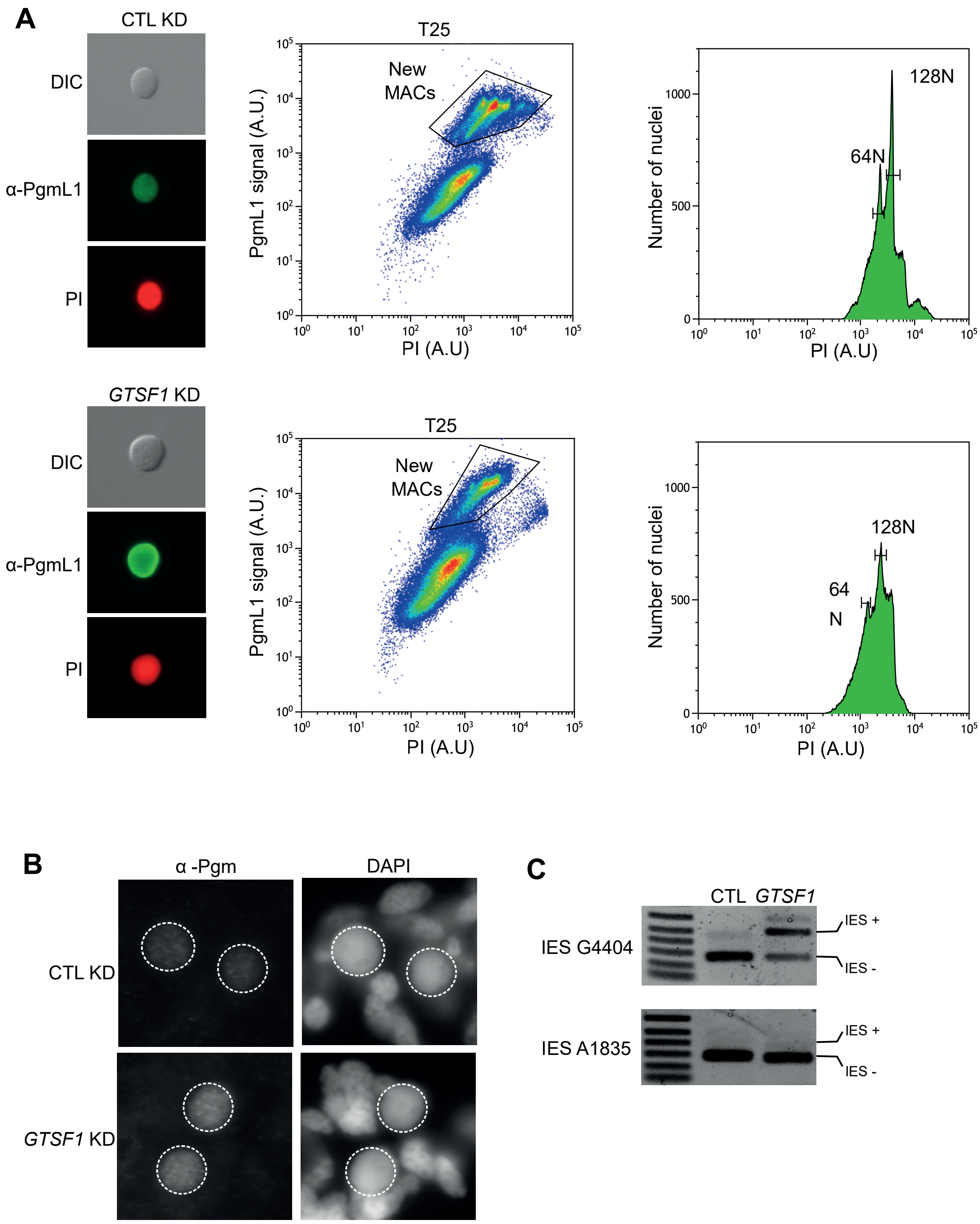

**Figure S3 - New developping MAC sorting by flow cytometry, Related to Figure 4**

A. Flow cytometry sorting of α-PgmL1 immunostained nuclei (Zangarelli et al., 2022) from control *ND7* (CTL) and *GTSF1* KD at T=25 hours after the onset of autogamy (DEV3 according to Zangarelli et al., 2022) (see Figure S1 for cytology). Left panels: images of the sorted nuclei in phase contrast (DIC), α-PgmL1 labeling and propidium iodide (PI) DNA labeling. Middle panels: plots of PgmL1 fluorescence intensity (y-axis; arbitrary units in log scale) versus PI fluorescence intensity (x-axis). New MACs gating used for nuclei sorting is indicated. Right panels: Histograms of PI-stained nuclei gated in the middle panels. The estimated ploidy level for most prominent peaks is shown.

B. Pgm immunostaining (α-Pgm2659-GP, Dubois et al., 2017) at T=25 hours after the onset of autogamy in control *ND7* (CTL) and *GTSF1* KD. New MACs and fragments of the maternal MAC are stained with DAPI, developing MACs are surrounded by a white dotted circle. The Pgm excision complex localizes in the new MAC upon *GTSF1* KD.

C. Enrichment for germline-specific sequences was checked by PCR around two IES sequences on DNA isolated from sorted new MACs upon control *ND7* (CTL) and *GTSF1* KD. Excised IES form (IES-) and non-excised form (IES+) are indicated.

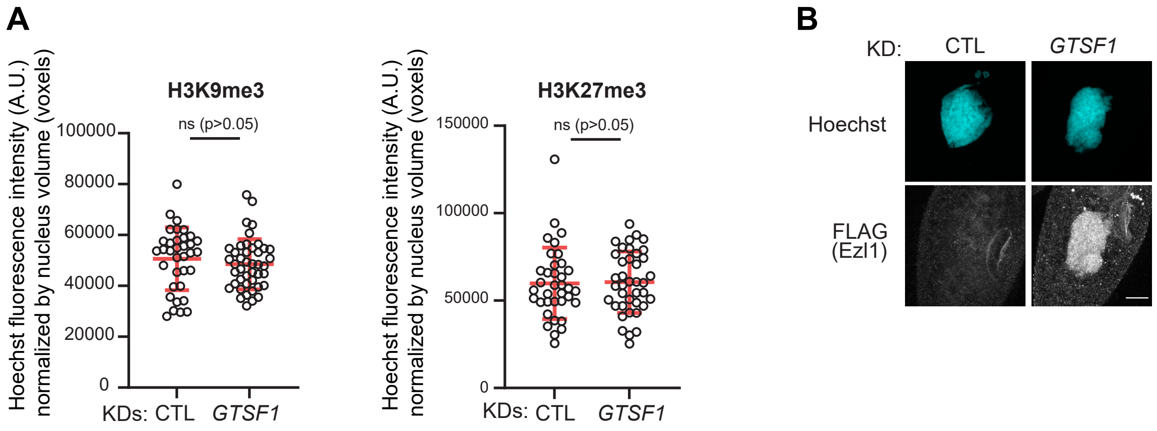

**Figure S4. Estimated size of new developing MAC. Gtsf1 controls Ezl1 levels. Related to Figure 5**

A. Boxplot of estimated nucleus (developing MAC) volume in voxels from the same data as in Figure 5A (T=15 hours). Number of nuclei > 30 in each condition. Estimation of nuclear volume indicated that *CTL* and *GTSF1* KD cell populations were at comparable stages. Bars correspond to mean +/- SD. Mann-Withney statistical test. n.s: non significant

B. FLAG immunostaining of cells expressing a *3XFLAG-HA-EZL1* functional transgene at T=0 hours after the onset of autogamy in *ICL7* (CTL) or *GTSF1* KDs. Scale bar, 10 µm.

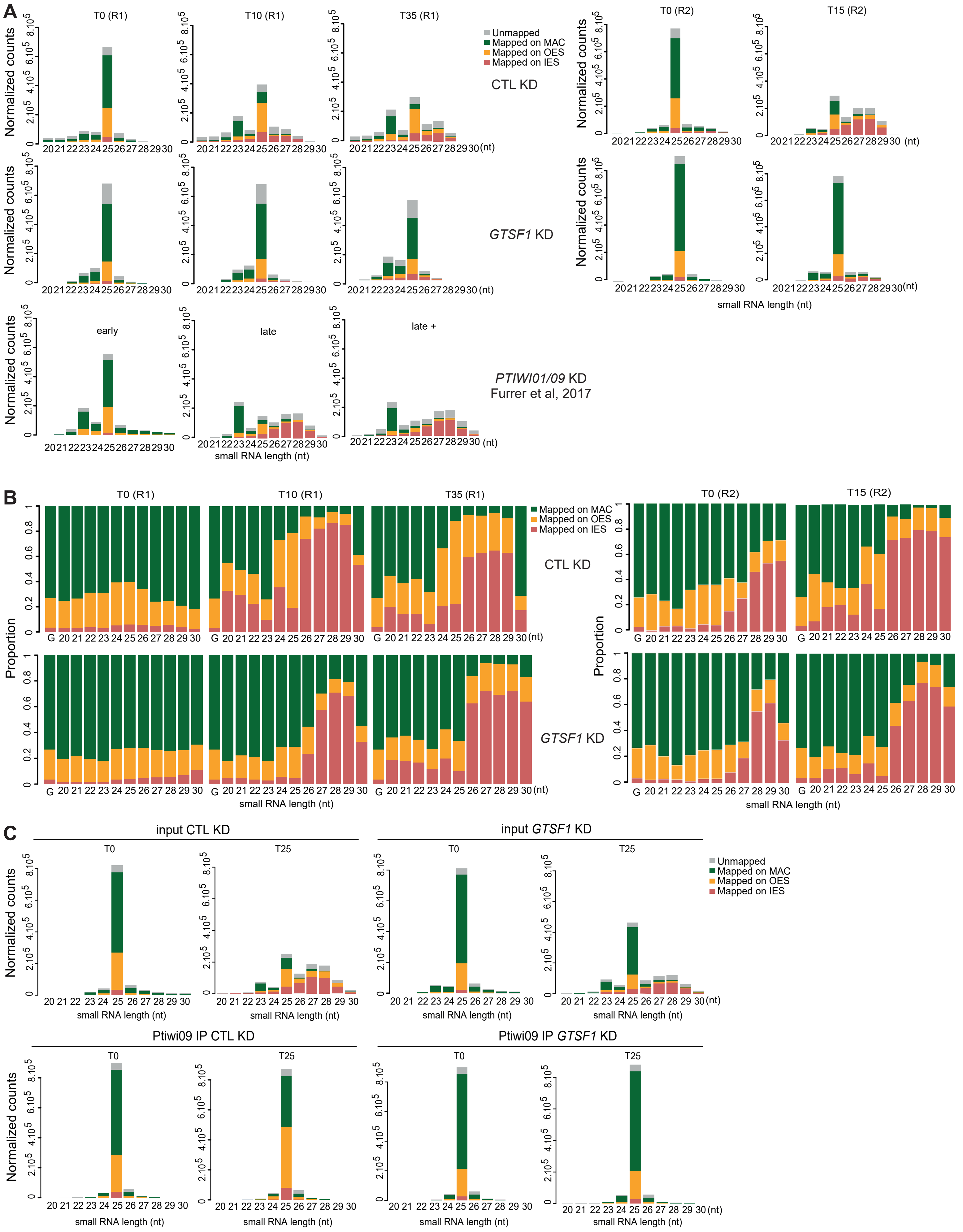

**Figure S5. Gtsf1 is required for scnRNA selection, Related to Figure 6**

Analysis of sRNA populations at different times of autogamy.

A. Bar plots show the normalized counts that map the MAC genome, IESs, or OES, at different time points of autogamy upon CTL and *GTSF1* KDs (2 replicates for each: R1 and R2).

B. Bar plots show the proportion of reads for each sample that map the MAC genome, IESs, or OES, at different time points of autogamy upon CTL and *GTSF1* KDs (2 replicates for each: R1 and R2).

C. Bar plots show the normalized counts for each sample before and after Ptiwi09 IP in *ND7* (CTL) or *GTSF1* KD that map the MAC genome, IESs, or OES, at 2 time points of autogamy upon CTL and *GTSF1* KD.

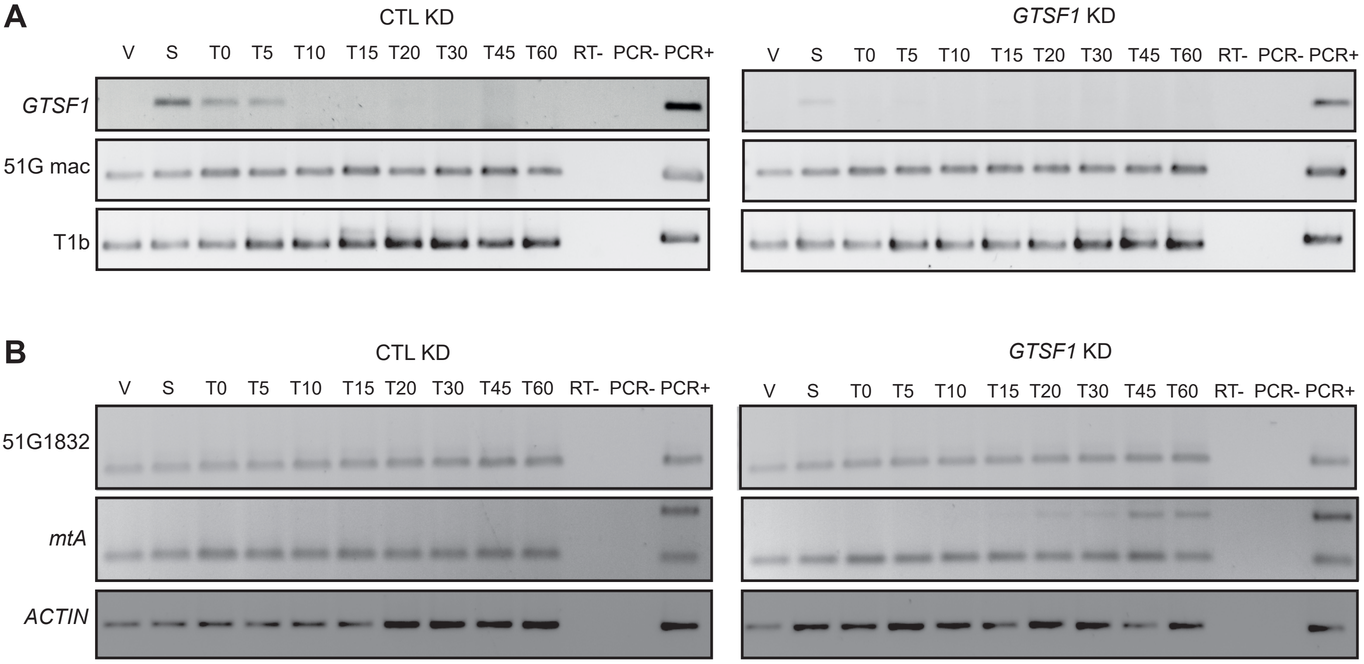

**Figure S6. Non coding maternal transcription is not affected upon *GTSF1* KD.**

Total RNAs, extracted at each time point, were reverse-transcribed and cDNAs were amplified by PCR with gene-specific primers. Loading controls are the constitutively-expressed genes encoding a trichocyst matrix protein (T1b) or actin. Total genomic DNA was used as a positive PCR control. Non-protein-coding RNAs are detected for the G surface antigen gene (51G mac) both during vegetative growth and during sexual events (Lepere et al., 2008). A. Detection of *GTSF1* and 51G mac ncRNA transcripts by RT-PCR in *ND7* (CTL) and *GTSF1* KD conditions. B. Detection of 51G1832 and *mtA* ncRNA transcripts by RT-PCR in *ND7* (CTL) and *GTSF1* KD conditions. Note the presence of an IES+ form for mtA, which is retained in *GTSF1* KD.

| | Name | Reference | MW (kDa) | Unique peptide | $Log^2$ FC | -Log10 p-value |
|---|---|---|---|---|---|---|
| Ptiwi09 IP | Caf1 | PTET.51.1.P0780031 | 45.62 | 8 | 13.06 | 1.85 |
| | Eap1 | PTET.51.1.P1310069 | 20.10 | 3 | 5.42 | 3.14 |
| | Eed | PTET.51.1.P0240079 | 43.92 | 3 | 16.19 | 4.81 |
| | Ezl1 | PTET.51.1.P1740049 | 69.69 | 6 | 11.38 | 1.36 |
| | Gtsf1 | PTET.51.1.P0490019 | 18.67 | 2 | 3.91 | 2.97 |
| | Pdsg1 | PTET.51.1.P0300085 | 37.98 | 3 | 9.56 | 1.32 |
| | PTETG3921 | PTET.51.1.P0390035 | 48.21 | 2 | 13.60 | 3.93 |
| | Ptiwi03 | PTET.51.1.P0030302 | 89.92 | 2 | 17.44 | 2.46 |
| | Ptiwi09 (bait) | PTET.51.1.P0660118 | 87.65 | 11 | 3.92 | 1.36 |
| | Rf2 | PTET.51.1.P1190062 | 74.74 | 9 | 13.02 | 1.72 |
| | Rf4 | PTET.51.1.P0570234 | 62.62 | 13 | 7.62 | 3.87 |
| | Rpb2 | PTET.51.1.P0480005 | 139.59 | 3 | 13.07 | 3.83 |
| Gtsf1 IP | Caf1 | PTET.51.1.P0780031 | 45.62 | 25 | 5.04 | 2.37 |
| | Eap1 | PTET.51.1.P1310069 | 20.10 | 5 | 5.52 | 2.02 |
| | Eed | PTET.51.1.P0240079 | 43.92 | 18 | 4.48 | 2.11 |
| | Ema1b | PTET.51.1.P0080253 | 174.48 | 30 | 3.68 | 1.49 |
| | Ezl1 | PTET.51.1.P1740049 | 69.69 | 30 | 4.88 | 2.08 |
| | Gtsf1 (bait) | PTET.51.1.P0490019 | 18.67 | 11 | 9.48 | 1.55 |
| | P0480005 | PTET.51.1.P0480005 | 139.59 | 18 | 2.06 | 1.55 |
| | P0850169 | PTET.51.1.P0850169 | 57.84 | 2 | 6.94 | 4.78 |
| | P1010056 | PTET.51.1.P1010056 | 40.59 | 2 | 5.79 | 1.49 |
| | Pdsg1 | PTET.51.1.P0300085 | 37.98 | 23 | 6.35 | 2.88 |
| | Ptiwi01 | PTET.51.1.P0710112 | 87.72 | 8 | 7.57 | 1.34 |
| | Ptiwi03 | PTET.51.1.P0030302 | 89.92 | 12 | 7.71 | 2.49 |
| | Ptiwi09 | PTET.51.1.P0660118 | 87.65 | 10 | 8.68 | 1.98 |
| | Rf2 | PTET.51.1.P1190062 | 74.74 | 48 | 5.66 | 1.92 |
| | Rf4 | PTET.51.1.P0570234 | 62.62 | 40 | 5.27 | 1.99 |
| | Rpb3a | PTET.51.1.P0510184 | 35.29 | 18 | 2.27 | 1.50 |
| | Rpb3b | GSPATG00038663001 | 20.08 | 10 | 2.35 | 1.34 |
| | Suz12.like | PTET.51.1.P0190277 | 34.65 | 17 | 4.48 | 2.07 |

**Table S1. Ptiwi09 and Gtsf1 interact together and with PRC2, Related to Figures 1 and 2.**
Top differential proteins in Flag IP compared to control IP for Ptiwi09 and for Gtsf1 IPs. A p-value inferior than 0.05 and a fold change superior than 2 were used to filter differential significant candidates. -Log10(p-value) > 1.3 is required to get a p-value < 0.05. Only proteins encoding genes with the same expression profiles than Ptiwi09 and Gtsf1 (early peak) were selected.

| Experiment n° | Transgene | KD gene | sexual progeny | |
|---|---|---|---|---|
| 1 - Production of sexual progeny (Figure 2C) | - | *ICL7* | Alive | 30 |
| | | | Dead | 0 |
| | | *GTSF1#1* | Alive | 0 |
| | | | Dead | 30 |
| | | *ICL7* | Alive | 30 |
| | | | Dead | 0 |
| | | *GTSF1#1* | Alive | 0 |
| | | | Dead | 30 |
| 2 - Production of sexual progeny (Figure 2C) | - | - | Alive | 29 |
| | | | Dead | 1 |
| | | *GTSF1#1* | Alive | 0 |
| | | | Dead | 30 |
| 3 - Production of sexual progeny (Figure 2C), DNA seq (Figure 4, S2), RNA-seq (Figure 4, S1 and S2) and small RNA seq (Figure 6B, S1 and S5) upon *GTSF1* KD | - | *ICL7* | Alive | 29 |
| | | | Dead | 1 |
| | | *GTSF1#1* | Alive | 0 |
| | | | Dead | 30 |
| 4 - Ezl1 localization by immunofluorescence (Figure S4), production of sexual progeny (Figure 2C) | - | *ICL7* | Alive | 13 |
| | | | Dead | 2 |
| | | *GTSF1#1* | Alive | 0 |
| | | | Dead | 15 |
| | *3XFLAG-HA-EZL1* | *ICL7* | Alive | 15 |
| | | | Dead | 0 |
| | | *GTSF1#1* | Alive | 3 |
| | | | Dead | 12 |
| | *3XFLAG-HA-EZL1* | *ICL7* | Alive | 15 |
| | | | Dead | 0 |
| | | *GTSF1#1* | Alive | 2 |
| | | | Dead | 13 |
| 5 - Gtsf1 genetic complementation experiments (Figure 2C), production of sexual progeny (Figure 2C), and localization at different stages of the sexual cycle (Figure 3) | - | *ICL7* | Alive | 30 |
| | | | Dead | 0 |
| | | *GTSF1#1* | Alive | 1 |
| | | | Dead | 29 |
| | *3XFLAG-HA-GTSF1* | *ICL7* | Alive | 28 |
| | | | Dead | 2 |
| | | *GTSF1#1* | Alive | 29 |
| | | | Dead | 1 |
| | *3XFLAG-HA-GTSF1* | *ICL7* | Alive | 28 |
| | | | Dead | 2 |
| | | *GTSF1#1* | Alive | 29 |
| | | | Dead | 1 |

| | | | | |
|---|---|---|---|---|
| | - | *ICL7* | Alive | 29 |
| | | | Dead | 1 |
| | | *GTSF1#1* | Alive | 0 |
| | | | Dead | 30 |
| 6 - Gtsf1 IP (Figure 2D, 2E, 2F and 2G), production of sexual progeny (Figure 2C) | *3XFLAG-HA-GTSF1* | *ICL7* | Alive | 29 |
| | | | Dead | 1 |
| | | *GTSF1#1* | Alive | 14 |
| | | | Dead | 1 |
| | *3XFLAG-HA-GTSF1* | *ICL7* | Alive | 15 |
| | | | Dead | 0 |
| | | *GTSF1#1* | Alive | 30 |
| | | | Dead | 0 |
| 7 - Ptiwi09 localization (Figure 7A) | *3XFLAG-HA-PTIWI09* | *ICL7* | Alive | 20 |
| | | | Dead | 7 |
| | | *GTSF1#1* | Alive | 30 |
| | | | Dead | 27 |
| | *3XFLAG-HA-PTIWI09* | *ICL7* | Alive | 30 |
| | | | Dead | 0 |
| | | *GTSF1#1* | Alive | 2 |
| | | | Dead | 28 |
| 8 - Production of sexual progeny (Figure 2C) | - | *ICL7* | Alive | 23 |
| | | | Dead | 7 |
| | | *GTSF1#1* | Alive | 0 |
| | | | Dead | 30 |
| 9 - Production of sexual progeny (Figure 2C) | - | *ICL7* | Alive | 26 |
| | | | Dead | 4 |
| | | *GTSF1#1* | Alive | 0 |
| | | | Dead | 30 |
| | | *ICL7* | Alive | 28 |
| | | | Dead | 2 |
| | | *GTSF1#1* | Alive | 2 |
| | | | Dead | 25 |
| 10 - Production of sexual progeny (Figure 2C) | — | *ICL7* | Alive | 48 |
| | | | Dead | 0 |
| | | *GTSF1#2* | Alive | 0 |
| | | | Dead | 48 |
| | | *ICL7* | Alive | 47 |
| | | | Dead | 1 |
| | | *GTSF1#2* | Alive | 0 |
| | | | Dead | 48 |
| 11 - Production of sexual progeny (Figure 2C) | — | *ICL7* | Alive | 46 |
| | | | Dead | 2 |
| | | *GTSF1#2* | Alive | 0 |
| | | | Dead | 48 |
| | | *ICL7* | Alive | 44 |
| | | | Dead | 4 |
| | | *GTSF1#2* | Alive | 2 |
| | | | Dead | 46 |
| | 3xFLAG-PTIWI09 | — | Alive | 48 |
| | | | Dead | 0 |

| | | | | |
|---|---|---|---|---|
| 12 - Ptiwi09 IP (Figure 1), production of sexual progeny (Figure 2C) | 3xFLAG-PTIWI09 | — | Alive | 47 |
| | | | Dead | 1 |
| | 3xFLAG-PTIWI09 | — | Alive | 46 |
| | | | Dead | 2 |
| | 3xFLAG-PTIWI09 | — | Alive | 48 |
| | | | Dead | 0 |
| | | — | Alive | 48 |
| | | | Dead | 0 |
| | | — | Alive | 48 |
| | | | Dead | 0 |
| 13 - Autogamy time course sRNA (Figure 6A, S1) ; sRNA seq (Figure S5) ; DNA (Figure 4 and S2) ; RT-PCR (Figure S6) upon *GTSF1* KD, production of sexual progeny (Figure 2C) | — | *ND7* | Alive | 47 |
| | | | Dead | 1 |
| | | *GTSF1#2* | Alive | 3 |
| | | | Dead | 45 |
| 14 -GTSF1-sorted DNA seq (Figure 4, S2 and S3), production of sexual progeny (Figure 2C) | — | *ND7* | Alive | 47 |
| | | | Dead | 1 |
| | | *GTSF1#2* | Alive | 2 |
| | | | Dead | 46 |
| 15 - Ptiwi09 RNA IP (Figure 6, S3), production of sexual progeny (Figure 2C) | 3xFLAG-PTIWI09 | *ICL7* | Alive | 48 |
| | | | Dead | 0 |
| | | *GTSF1#2* | Alive | 2 |
| | | | Dead | 46 |
| | — | *ICL7* | Alive | 48 |
| | | | Dead | 0 |
| | | *GTSF1#2* | Alive | 0 |
| | | | Dead | 48 |
| 16 - Autogamy time course (Figure 7B), production of sexual progeny (Figure 2C) | 3xFLAG-PTIWI09 | *L4440* | Alive | 48 |
| | | | Dead | 0 |
| | | *GTSF1#2* | Alive | 0 |
| | | | Dead | 48 |
| | — | *L4440* | Alive | 45 |
| | | | Dead | 3 |
| | | *GTSF1#2* | Alive | 0 |
| | | | Dead | 48 |
| 17 - Ptiwi09 IP (Figure 7C), production of sexual progeny (Figure 2C) | 3xFLAG-PTIWI09 | *ICL7* | Alive | 48 |
| | | | Dead | 0 |
| | | *GTSF1#2* | Alive | 1 |
| | | | Dead | 47 |
| | — | *ICL7* | Alive | 48 |
| | | | Dead | 0 |
| | | *GTSF1#2* | Alive | 2 |
| | | | Dead | 46 |
| 18 - PGM staining (Figure S3), production of sexual progeny (Figure 2C) | — | *ND7* | Alive | 48 |
| | | | Dead | 0 |
| | | *GTSF1#2* | Alive | 1 |
| | | | Dead | 47 |
| 19 - DNA seq upon *Ptiwi01/09* KD (R1) (Figure S2) | — | *PTIWI01/09* | Alive | |
| | | | Dead | 63% |

**Table S2. Production of sexual progeny following RNAi-mediated gene silencing**
In each different experiment, the number of cells that survived or died is indicated.

| Sequencing | Sample | Label | ENA Accession | Reference | Number of reads | Aligned reads on the MAC | | Aligned reads on the MIC | |
|---|---|---|---|---|---|---|---|---|---|
| DNAseq | KLEB | MAC | ERS452529 | Lhuillier-Akakpo et al. 2014 | 106 056 122 | 104 443 929 | 98% | 105 570 391 | 100% |
| DNAseq | PTET_ND7_RNAi_T25_AlgFACS_JkN_REGN50 | ND7_Alg | ERS16327826 | this study | 38 321 642 | 35 879 076 | 94% | 38 065 563 | 99% |
| DNAseq | PTET_GTSF1L-RNAi_TotalDNA_d4_JKN | GTSF1L_R2 | ERS16327827 | this study | 121 834 466 | 116 955 124 | 96% | 121 634 908 | 100% |
| DNAseq | PTET_ZF1_RNAi_gDNA_T50_DUHA-166 | ZF1_R1 | ERS16327828 | this study | 58 868 356 | 55 622 433 | 94% | 58 332 259 | 99% |
| DNAseq | PTET_GTSF1L_RNAi_T25_AlgFACS_JkN_REGN51 | GTSF1L_Alg | ERS16327829 | this study | 58 418 092 | 47 635 163 | 82% | 57 949 376 | 99% |
| DNAseq | MicGSC_BCP_AAIOSF_2_HiSeq | MIC | ERX4616645 | Sellis et al. 2021 | 181 407 606 | 153 120 470 | 84% | 179 098 715 | 99% |
| DNAseq | PGM-1_FACS_ANLG | PGM | SAMN05323661 | Guérin et al, 2017 | 115 558 914 | 101 303 406 | 88% | 115 259 143 | 100% |
| DNAseq | Ezl174-1_RNAi_r1_r2 | EZL_r1 | ERX466733 | Lhuillier-Akakpo et al. 2014 | 99 695 690 | 87 432 783 | 88% | 97 164 770 | 97% |
| DNAseq | Ezl174-2_RNAi_r1 | EZL_r2 | ERS452532 | Lhuillier-Akakpo et al. 2014 | 92 725 940 | 82 152 917 | 89% | 92 446 907 | 100% |
| DNAseq | PTIWI19_RNAi_r1 | PTIWI19_r1 | ERS16327830 | this study | 49 410 804 | 46 824 351 | 95% | 48 884 765 | 99% |
| DNAseq | PTET_PTIWI_1_9_KD_DNA_ERR1918503 | PTIWI19_r2 | ERS1656549 | Furrer et al. 2017 | 118 338 520 | 108 845 895 | 92% | 117 994 275 | 100% |
| DNAseq | DCL2_3_RNAi_r1_HBJ-1 | DCL2_3_r1 | PRJNA184719 | Sandoval et al. 2014 | 100 445 166 | 89 354 921 | 89% | 99 646 333 | 99% |
| DNAseq | Dcl2-3_RNAi_r2 | DCL2_3_r2 | ERX466736 | Lhuillier-Akakpo et al. 2014 | 99 744 888 | 92 915 718 | 93% | 99 600 735 | 100% |
| DNAseq | DCL5_RNAi_r1_HBJ-2 | DCL5 | PRJNA184719 | Sandoval et al. 2014 | 90 105 744 | 88 212 339 | 98% | 89 877 717 | 100% |
| DNAseq | DCL235_KD_MaN | DCL235 | SAMEA3726521 | Swart et al. 2017 | 54 294 800 | 51 349 502 | 95% | 53 581 515 | 99% |
| mRNAseq | ICL7_T0_RNA_DUHA140 | ICL7_T0 | ERS6679030 | Miro-Pina et al. 2022 | 79765916 | 78958586 | 99% | 77571546 | 97% |
| mRNAseq | ICL7_T10_RNA_DUHA141 | ICL7_T10 | ERS6679031 | Miro-Pina et al. 2022 | 86517736 | 85652820 | 99% | 84056363 | 97% |
| mRNAseq | ICL7_T35_RNA_DUHA142 | ICL7_T35 | ERS6679032 | Miro-Pina et al. 2022 | 109355820 | 108387011 | 99% | 106333998 | 97% |
| mRNAseq | ICL7_T50_RNA_DUHA143 | ICL7_T50 | ERS6679033 | Miro-Pina et al. 2022 | 100914984 | 99997163 | 99% | 98007556 | 97% |
| mRNAseq | PGM-T2_mRNA_CACTCA | PGM_T2 | ERS14842492 | Bazin-Gélis et al. 2023 | 81069812 | 77132399 | 95% | 74844554 | 92% |
| mRNAseq | PGM-T10_mRNA_CTCAGA | PGM_T10 | ERS14842490 | Bazin-Gélis et al. 2023 | 71443788 | 70294491 | 98% | 68207123 | 95% |
| mRNAseq | PGM-T30_mRNA_ATTCCT | PGM_T30 | ERS14842493 | Bazin-Gélis et al. 2023 | 80832994 | 75672689 | 94% | 73986877 | 92% |
| mRNAseq | PGM-T40_mRNA_CACGAT | PGM_T40 | ERS14842494 | Bazin-Gélis et al. 2023 | 79637140 | 77396584 | 97% | 76217304 | 96% |
| mRNAseq | PTET_RNAs_ZF1-T0_DUHA161 | ZF1_T0 | ERS16327831 | this study | 123476646 | 121939263 | 99% | 119767868 | 97% |
| mRNAseq | PTET_RNAs_ZF1-T10_DUHA162 | ZF1_T10 | ERS16327832 | this study | 118472868 | 116661852 | 98% | 114875670 | 97% |
| mRNAseq | PTET_RNAs_ZF1-T35_DUHA163 | ZF1_T35 | ERS16327833 | this study | 122393108 | 120117010 | 98% | 119019613 | 97% |
| mRNAseq | PTET_RNAs_ZF1-T50_DUHA164 | ZF1_T50 | ERS16327834 | this study | 124817826 | 122582514 | 98% | 121512408 | 97% |
| mRNAseq | EZL1_T0_RNA_DUHA144 | EZL1_T0 | ERS6679026 | Miro-Pina et al. 2022 | 72271092 | 71578739 | 99% | 70385457 | 97% |
| mRNAseq | EZL1_T10_RNA_DUHA145 | EZL1_T10 | ERS6679027 | Miro-Pina et al. 2022 | 69918912 | 68772590 | 98% | 68009496 | 97% |
| mRNAseq | EZL1_T35_RNA_DUHA146 | EZL1_T35 | ERS6679028 | Miro-Pina et al. 2022 | 107514064 | 99201883 | 92% | 104563941 | 97% |
| mRNAseq | EZL1_T50_RNA_DUHA147 | EZL1_T50 | ERS6679029 | Miro-Pina et al. 2022 | 95433848 | 86764000 | 91% | 92662006 | 97% |
| mRNAseq | ARN_PTIWI1-9_71a-T0_S4 (DUHA90) | PTIWI1_9_T0 | ERS6678315 | Miro-Pina et al. 2022 | 105639316 | 104163879 | 99% | 102551416 | 97% |
| mRNAseq | ARN_PTIWI1-9_71a-T10_S5 (DUHA91) | PTIWI1_9_T10 | ERS6678316 | Miro-Pina et al. 2022 | 85631710 | 84289504 | 98% | 83276546 | 97% |
| mRNAseq | ARN_PTIWI1-9_71a-T35_S6 (DUHA92) | PTIWI1_9_T35 | ERS6678317 | Miro-Pina et al. 2022 | 97923744 | 95405812 | 97% | 95157309 | 97% |
| mRNAseq | ARN_PTIWI1-9_71a-T50_S7 (DUHA93) | PTIWI1_9_T50 | ERS6678318 | Miro-Pina et al. 2022 | 97927868 | 93022880 | 95% | 95185568 | 97% |

| | | | | | | | | | |
|---|---|---|---|---|---|---|---|---|---|
| sRNAseq | ICL7T0_ATCACG_L006_007_R1_001 | ICL7 T0_R1 | PRJEB46608 | Miro-Pina et al. 2022 | 14640580 | 7232735 | 49% | 11 496 959 | 79% |
| sRNAseq | ICL7T10_CGATGT_L006_007_R1_001 | ICL7 T10_R1 | ERS14549878 | Miro-Pina et al. 2023 | 21656218 | 4800906 | 22% | 14 886 100 | 69% |
| sRNAseq | ICL7T35_TGACCA_L006_007_R1_001 | ICL7 T35_R1 | ERS14549879 | Miro-Pina et al. 2023 | 5810542 | 1178480 | 20% | 3 969 579 | 68% |
| sRNAseq | sRNA_Control_GTSF1L_T0_JKN | CTL T0_R2 | ERS16327835 | this study | 7470729 | 3977980 | 53% | 6 349 560 | 85% |
| sRNAseq | sRNA_Control_GTSF1L_T15_JKN | CTL T15_R2 | ERS16327836 | this study | 6506248 | 1114163 | 17% | 5 162 530 | 79% |
| sRNAseq | sRNA_GTSF1L_T0_JKN | GTSF1L T0_R2 | ERS16327837 | this study | 6941745 | 4685083 | 67% | 6 259 542 | 90% |
| sRNAseq | sRNA_GTSF1L_T15_JKN | GTSF1L T15_R2 | ERS16327838 | this study | 5915826 | 3488112 | 59% | 5 215 627 | 88% |
| sRNAseq | sRNAs_ZF1_T0_DUHA184 | ZF1 T0_R1 | ERS16327839 | this study | 22759952 | 11888719 | 52% | 16 188 772 | 71% |
| sRNAseq | sRNAs_ZF1_T10_DUHA185 | ZF1 T10_R1 | ERS16327840 | this study | 26350856 | 13329934 | 51% | 18 727 289 | 71% |
| sRNAseq | sRNAs_ZF1_T35_DUHA186 | ZF1 T35_R1 | ERS16327841 | this study | 18305725 | 7956240 | 43% | 12 645 019 | 69% |
| sRNAseq | sRNA_ICL-RNAi_Input_T0_JkN_S149 | Input T0 CTL | ERS16327842 | this study | 9081430 | 5639683 | 62% | 8 216 558 | 90% |
| sRNAseq | sRNA_ICL-RNAi_Input_T25_JkN_S150 | Input T25 CTL | ERS16327843 | this study | 10607853 | 1641339 | 15% | 8 777 369 | 83% |
| sRNAseq | sRNA_ICL-RNAi_IP-PTIWI09_T0_JkN_S151 | IPptiwi T0 CTL | ERS16327844 | this study | 11943676 | 7355448 | 62% | 10 980 105 | 92% |
| sRNAseq | sRNA_ICL-RNAi_IP-PTIWI09_T25_JkN_S152 | IPptiwi T25 CTL | ERS16327845 | this study | 7947370 | 3023379 | 38% | 7 265 026 | 91% |
| sRNAseq | sRNA_GTSF1L-RNAi_Input_T0_JkN_S153 | Input T0 GTSF1L | ERS16327846 | this study | 9063379 | 6263529 | 69% | 8 226 346 | 91% |
| sRNAseq | sRNA_GTSF1L-RNAi_Input_T25_JkN_S154 | Input T25 GTSF1L | ERS16327847 | this study | 8426310 | 3470089 | 41% | 6 949 269 | 82% |
| sRNAseq | sRNA_GTSF1L-RNAi_IP-PTIWI09_T0_JkN_S155 | IPptiwi T0 GTSF1L | ERS16327848 | this study | 10815354 | 7522230 | 70% | 9 961 101 | 92% |
| sRNAseq | sRNA_GTSF1L-RNAi_IP-PTIWI09_T25_JkN_S156 | IPptiwi T25 GTSF1L | ERS16327849 | this study | 7531389 | 5276938 | 70% | 6 930 118 | 92% |

**Table S3. Sequencing data and mapping statistics**

DNA-seq, RNA-seq and sRNA-seq data were used in this study. For each sequencing sample the ENA acession is specified, followed by the number of reads sequenced and the mapped reads on the MAC and MIC reference.

| Name | Sequence (5' to 3') | Locus (ID) | Source | Application |
|---|---|---|---|---|
| Actin_qPCR_for | TGAAGCTCCAATGAATCCAA | Actin 1-1 (PTET.51.1.G0130204) | Frapporti et al., 2019 | ChIP-qPCR |
| Actin_qPCR-rev | TCCTGAAGCATAGAGTGAGA | Actin 1-1 (PTET.51.1.G0130204) | Frapporti et al., 2019 | ChIP-qPCR |
| GAPDH_qPCR_F2 | ATTTTGGTATTGTTGAGGGT | GAPDH (PTET.51.1.G0380195) | Frapporti et al., 2019 | ChIP-qPCR |
| GAPDH_qPCR_R2 | CTCCAGTCTTTTCCACCTTT | GAPDH (PTET.51.1.G0380195) | Frapporti et al., 2019 | ChIP-qPCR |
| Anchois.173_F2 | TTCCAAGCTGATTCGATTATTAAA | Anchois B (IESPGM.PTET51.1.173.70900) | Frapporti et al., 2019 | ChIP-qPCR |
| Anchois.173_R2 | ACTTCTTGTTTCATTTGTTAGACT | Anchois B (IESPGM.PTET51.1.173.70900) | Frapporti et al., 2019 | ChIP-qPCR |
| Oligo #551 (RT31010) | ACAAGATTGACCAGGACTTATT | RT31010 (ms4410_NODE_3768_length_13900 _cov_21.582806_RT31010_Group4_nonLTR:ClassI:LINE) | Frapporti et al., 2019 | ChIP-qPCR |
| Oligo #552 (RT31010) | ATATCATTCACTCCTGCAATCT | RT31010 (ms4410_NODE_3768_length_13900 _cov_21.582806_RT31010_Group4_nonLTR:ClassI:LINE) | Frapporti et al., 2019 | ChIP-qPCR |
| Oligo #559 (RT42890) | TTAATTGAAGGCGAAGAAAGAC | RT42890 (ms1831_NODE_10132_length_49470 _cov_21.140064_RT42890_Group4_nonLTR:ClassI:LINE) | Frapporti et al., 2019 | ChIP-qPCR |
| Oligo #560 (RT42890) | TTAATTGAAGGCGAAGAAAGAC | RT42890 (ms1831_NODE_10132_length_49470 _cov_21.140064_RT42890_Group4_nonLTR:ClassI:LINE) | Frapporti et al., 2019 | ChIP-qPCR |
| Oligo #723 (RT48639-2) | ATCATCTTTCCCTCACATCG | RT48639-2 (ms4963_NODE_3562_length_10237 _cov_31.527792_RT48639exp_Group2_nonLTR:ClassI:LINE) | Frapporti et al., 2019 | ChIP-qPCR |
| Oligo #724 (RT48639-2) | AGATTTCACGCTTCAGTTCT | RT48639-2 (ms4963_NODE_3562_length_10237 _cov_31.527792_RT48639exp_Group2_nonLTR:ClassI:LINE) | Frapporti et al., 2019 | ChIP-qPCR |
| expGTSF1U | ATGTAATTAAAATTGAAATCCTACGAACTAG | GTSF1 (PTET.51.1.G0490019) | this study | RT-PCR |
| expGTSF1L | GTTTAATTCTTTTGACCGAGGACTC | GTSF1 (PTET.51.1.G0490019) | this study | RT-PCR |
| T1b-3' | TTGAGTTGGGATTTGACATAATCGGTGAA | T1b (PTET.51.1.G0980135) | Maliszewska-Olejniczak et al., 2015 | RT-PCR |
| T1b-5' (2) | TCTAATTAAACCAAGAACACGCTGAATTCC | T1b (PTET.51.1.G0980135) | Maliszewska-Olejniczak et al., 2015 | RT-PCR |
| 51G18 | ACTGTTGCTACACATTGTGCATATGTTACT | 51G mac | Maliszewska-Olejniczak et al., 2015 | RT-PCR |
| 51G17 | GATCAAGTCCAGTTCCTGTTATAGAACTAC | 51G mac | Maliszewska-Olejniczak et al., 2015 | RT-PCR |
| G1832 | GCTATAACTCTTGAAGCTGCTTGTAATATG | 51G | Lhuillier-Akakpo et al., 2014 | RT-PCR |
| G1832 | TTGTCAATGAGCCATTAACAGTTGCTGGAT | 51G | Lhuillier-Akakpo et al., 2014 | RT-PCR |
| ActinF | AGACCACCCAGCTCTTTTGA | Actin 1-1 (PTET.51.1.G0130204) | this study | RT-PCR |
| ActinR | TTGGGACTGTGTGTGAGACA | Actin 1-1 (PTET.51.1.G0130204) | this study | RT-PCR |
| mtApromF | CTTATTCTGCCTTCTCTTGAAATGC | mtA | this study | RT-PCR |
| mtApromR | AGGTCATCTCTTTCATTAAATTCCT | mtA | this study | RT-PCR |

**Table S4. Oligonucleotide sequences, Related to Figures 5 and S6**
